# THE OLFACTORY RECEPTOR Olfr78 REGULATES DIFFERENTIATION OF ENTEROCHROMAFFIN CELLS IN THE MOUSE COLON

**DOI:** 10.1101/2023.04.19.536389

**Authors:** Gilles Dinsart, Morgane Leprovots, Anne Lefort, Frédérick Libert, Yannick Quesnel, Alex Veithen, Gilbert Vassart, Sandra Huysseune, Marc Parmentier, Marie-Isabelle Garcia

## Abstract

The gastrointestinal epithelium constitutes a chemosensory system for microbiota-derived metabolites such as Short Chain Fatty Acids (SCFA). In this study, we investigated spatial distribution of Olfr78, one of the SCFA receptors, in the mouse intestine and studied the transcriptome of colon enteroendocrine cells expressing Olfr78. The receptor is principally detected in the enterochromaffin and L subtypes in the proximal and distal colon, respectively. Using the Olfr78-GFP and VilCre/Olfr78flox transgenic mouse lines, we reveal that loss of epithelial Olfr78 results in impaired enterochromaffin cell differentiation, blocking cells in an undefined secretory lineage state. This is accompanied by dysbiosis, characterized by an increased *Firmicutes*/*Bacteroidete*s ratio, as well as a less efficient antioxidant system in colon crypts. Using organoid cultures, we further show that maintenance of enterochromaffin cells involves activation of the Olfr78 receptor via the SCFA ligand acetate. Altogether, this work provides evidence that Olfr78 contributes to colon homeostasis by regulating enterochromaffin cell differentiation.

## INTRODUCTION

The gastrointestinal epithelium is an important contributor to endocrine physiology and metabolic control by constituting a chemosensory system for external toxic agents, as well as diet-derived nutrients or microbiota-derived metabolites. This “sensory” function is mainly exerted by a subset of epithelial cells, known as enteroendocrine cells (EEC). In response to stimuli, these highly specialized cells secrete neurotransmitters or a variety of hormones having paracrine digestive and endocrine functions (Gribble & Reimann, 2019). Recent reports have uncovered the extraordinary complexity of this lineage, composed of multiple subtypes, each producing a defined set of specific hormones/signaling molecules (Basak et al., 2017; Egerod et al., 2012; Haber et al., 2017; Habib et al., 2012; Roth et al., 1992). In addition, EEC subtypes are also differentially distributed along the digestive tract, according to both proximal-to-distal and crypt-villus axes (Beumer et al., 2018; Haber et al., 2017; Roth et al., 1992). Two main EEC subtypes populate the colon, the so-called L cells devoted to secretion of GLP-1 and PYY peptides and the enterochromaffin cell subtype (EC) that secretes most of the serotonin (5-HT for 5-hydroxytryptamine) produced by the body (Gribble & Reimann, 2019).

Till recently, not much was known about the identity of the molecules that sense the chemical stimuli enabling signal transduction in EECs to promote hormone or neurotransmitter secretion. Accumulated evidence revealed the key role of some G protein-coupled receptors (GPCR) in such process. These receptors recognize as natural ligands the luminal short chain fatty acids (SCFA) generated from microbiota-derived metabolites (Bellono et al., 2017; Pluznick, 2016). The most abundant SCFAs, acetate, propionate and butyrate, are particularly concentrated in the colon, where they reach the millimolar concentration range (Cong et al., 2022). The FFAR2/GPR43 and FFAR3/GPR41 receptors, expressed in the EEC L subtype, recognize acetate, propionate, butyrate and isovalerate as ligands (Audouze et al., 2014; Le Poul et al., 2003; Pluznick et al., 2013; Saito et al., 2009). Whether they are involved in hormonal secretion in these cells is still debated (Christiansen et al., 2018; Psichas et al., 2015; Tolhurst et al., 2012). Loss of function studies in mice have demonstrated that the mouse orthologs Ffar2/Gpr43 and Ffar3/Gpr41 receptors participate to intestinal homeostasis by regulating inflammation in response to chemically induced colitis (Kim et al., 2013). The two other SCFA binding molecules are the olfactory receptors OR51E1 and OR51E2 (the human orthologs of the mouse receptors Olfr558 and Olfr78, respectively). Although odorant receptors are mainly expressed in olfactory sensory neurons of the olfactory epithelium, Olfr78 and Olfr558 expression is also detected in other tissues including the digestive tract (Bellono et al., 2017; Billing et al., 2019; Fleischer et al., 2015; Lund et al., 2018). It has been demonstrated that binding of SCFAs (isovalerate, butyrate) to the mouse Olfr558 receptor in EC cells of the small intestine activates basolateral 5-HT release from secretory granules and thereby stimulates afferent nerve fibers (Bellono et al., 2017). The Olfr78 receptor (human ortholog OR51E2) that recognizes acetate and propionate as ligands, was firstly detected in the EEC L subtype (Fleischer et al., 2015). Moreover, transcriptome analyses have also reported its expression in EC cells in the colon (Lund et al., 2018). Using the transgenic knockin/knockout Olfr78-GFP/LacZ mouse line, Kotlo et al. have demonstrated that absence of Olfr78 expression is associated with higher levels of intestinal inflammation and worse histopathological score as compared to control mice in an experimental model of colitis (Kotlo et al., 2020). However, the role, if any, of Olfr78 in intestinal epithelium under homeostatic conditions remains to be addressed.

In the present work, we investigated the complete expression profile of Olfr78 in gut and uncovered the biological function of this SCFA receptor in mouse colon epithelium using transgenic mouse lines and organoid cultures. Our findings reveal that signaling through the Olfr78 receptor regulates EC lineage differentiation in colon epithelium and participates to tissue homeostasis.

## RESULTS

### SCFA receptors exhibit unique expression profiles along the small intestine and the colon

To fully dissect the distribution pattern of SCFA receptors, namely Olfr78, Olfr558, Ffar2 and Ffar3, along the mouse small and large intestines, we first analyzed their gene expression on mouse control tissues by qRT-PCR experiments. While *Ffar2* and *Ffar3* transcripts were detected at similar levels along the intestine, higher levels of *Olfr78* and *Olfr558* transcripts were found in the colon as compared to the small intestine (Figure 1A). *In situ* hybridization studies further showed that *Olfr78*, *Olfr558* and *Ffar3* were expressed in discrete countable cells in the epithelium (Figure 1B, C and D). In the proximal colon, epithelial *Olfr78* and *Olfr558*-expressing cells were mostly located at the crypt base, while *Ffar3*- positive (^+ve^) cells were mainly found at the top of the crypts, near the lumen, suggesting that Olfrs and Ffar3 receptors may not be co-expressed in the same cell types (Figure 1B, D). Regarding *Ffar2*, this receptor was more diffusely expressed and showed a decreasing gradient from the bottom to the bottom-half of colon crypts (Figure 1B). In addition, *Olfr78* and *Olfr558* were expressed in isolated mesenchymal cells and submucosal and myenteric plexuses in the colon (Figure EV1A, B). Of note, expression of *Olfr558* in the distal colon was only attributed to non-epithelial cells (Figures 1A-C and EV1B). Finally, in agreement with a previous report (Nøhr et al., 2013), *Ffar3*^+ve^ cells were present in the myenteric plexuses (from ileum to distal colon) and *Ffar2* expression was detected in submucosal cells (likely leukocytes) (Figure EV1C, D). Altogether, these data revealed a unique expression profile of SCFA receptors along the proximal-to-distal and crypt bottom-top villus axes in the gut.

**Figure 1.**
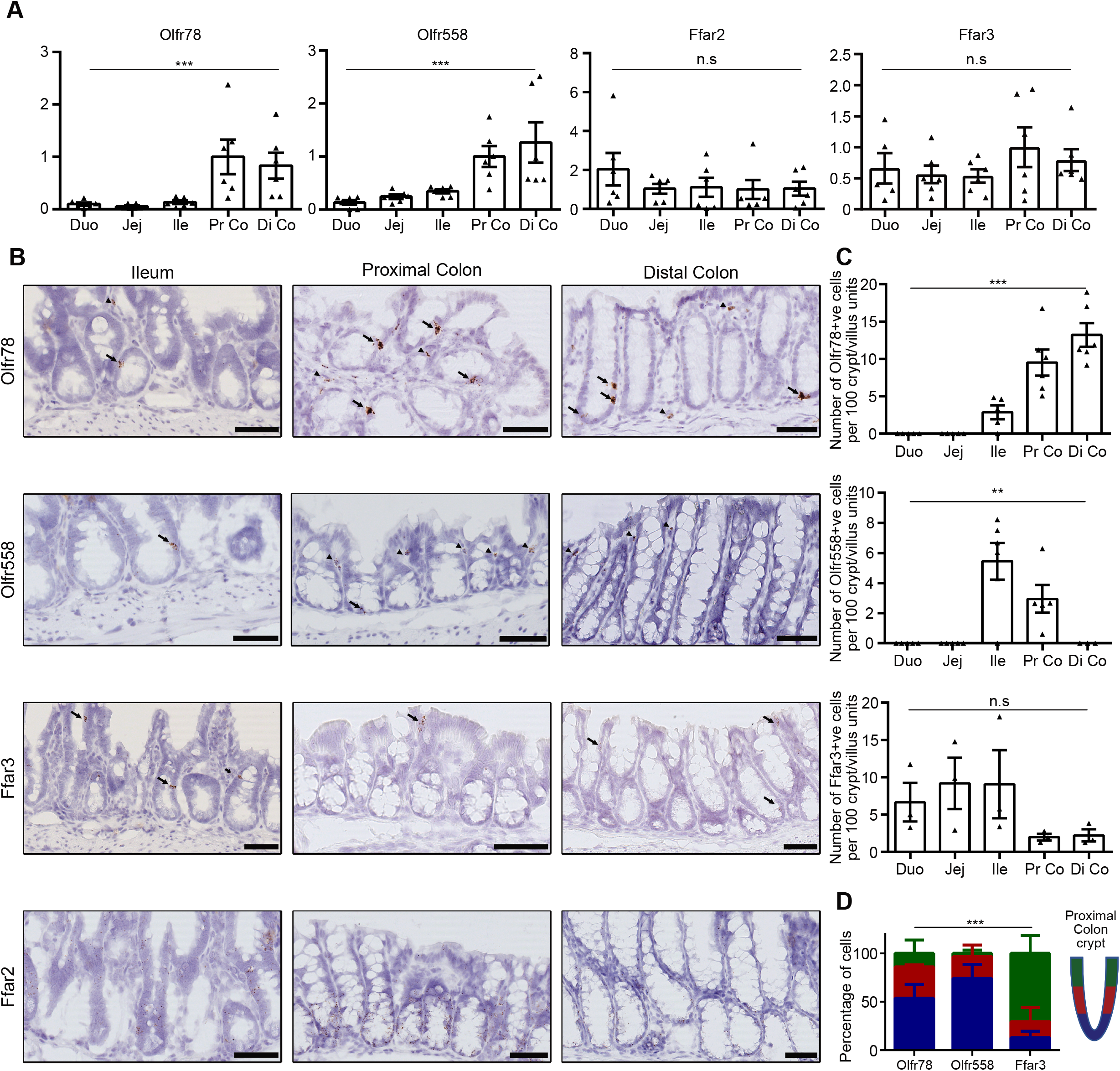
SCFA receptors exhibit unique expression profiles along the small intestine and colon. A. The expression profile of SCFA receptors was analyzed by qRT-PCR on wild-type (WT) mouse intestinal biopsies from duodenum (Duo), jejunum (Jej), ileum (Ile), proximal colon (Pr Co) and distal colon (Di Co). Relative expression levels were arbitrary set to 1 in Pr Co samples. Each symbol indicates the value for a given mouse (n=6). B. The expression profile of SCFA receptors was analyzed on WT mouse tissues by RNAscope. Arrows and arrowheads point to epithelial and non-epithelial expressing cells, respectively. C. Quantification of the number of cells expressing the Olfr78, Olfr558 and Ffar3 SCFA receptors per 100 crypt/villus sections from pictures obtained with RNAscope. Each symbol indicates the value of a given mouse (n=3 to 6). D. Distribution of Olfr78, Olfr558 and Ffar3 positive cells along proximal colon crypts (blue: bottom, red: middle, green: top of the crypt, n=3 mice). Data information: Scale bars = 50 µm (B). Data are represented as mean ± SEM. (A) Kruskal-Wallis tests: *** P=0.002 (Olfr78), *** P=0.002 (Olfr558) and n.s = not significant (Ffar3) with Dunn’s multiple comparison tests: * P=0.0370 Duo vs Pr Co and Duo vs Di Co, ** P=0.045 Jej vs Pr Co and Jej vs Di Co (Olfr78); ** P=0.027 Duo vs Pr Co, ** P=0.0016 Duo vs Di Co, * P=0.0351 Jej vs Di Co (Olfr558). (C) Kruskal-Wallis tests: *** P=0.001 (Olfr78), ** P=0.0013 (Olfr558) and n.s = not significant (Ffar3) with Dunn’s multiple comparison tests: * P=0.0370 Duo vs Pr Co and Duo vs Di Co, ** P=0.045 Jej vs Pr Co and Jej vs Di Co (Olfr78); ** P=0.027 Duo vs Pr Co, ** P=0.0016 Duo vs Di Co, * P=0.0351 Jej vs Di Co (Olfr558). (D)Two-way ANOVA test (interaction *** p=0.008) with Tukey’s multiple comparison tests: bottom crypts: ** p=0.0067 for Olfr558 vs Ffar3; top crypts: * p=0.017 for Olfr78 vs Ffar3 and ** p=0.0031 for Olfr558 vs Ffar3.

### Olfr78 is expressed in different subtypes of enteroendocrine cells in the colon

Since *Olfr78* expression was particularly abundant in the colon where SCFA ligands are predominant, we further explored its expression profile in colon epithelial cells using the knockin-knockout reporter Olfr78^tm1Mom^ mouse strain (thereafter referred as Olfr78-GFP). As depicted in Figure 2A, Olfr78-GFP mice harbor a GFP-IRES-tauLacZ cassette in place of the coding exon of *Olfr78* (Bozza et al., 2009). Using GFP reporter as a surrogate to identify Olfr78^+ve^ cells, Epcam^+ve^ /GFP^+ve^ cells were sorted from the colon of 4 heterozygous (HE) Olfr78^GFP/+^ mice (two pools, each obtained from 2 individual mice) by flow cytometry (Figures 2A and EV2A). All epithelial cells (Epcam^+ve^) were also isolated from a wild-type (WT) colon (Figure 2A). Following bulk RNA sequencing (seq) of sorted cells, we compared the transcriptome of epithelial Olfr78^+ve^ cells to that of all Epcam^+ve^ cells with the Degust software to identify significantly up and downregulated genes in Olfr78-expressing cells. Using a cut-off of false discovery rate (FDR) < 0.001 and a log2 fold-change of > 2 or < −2, we identified 905 enriched genes and 321 de-enriched genes (Figure 2B). As expected, *Olfr78* expression was 60-fold enriched in GFP^+ve^ cells (Figure 2B and 2C). Regarding other SCFA receptors, *Olfr558* and *Ffar2* were found enriched by 50- and 4-fold, respectively, whereas *Ffar3* expression was not detected at significant levels in GFP^+ve^ cells (Figure 2B, 2C). High levels of expression of general EEC (*Chga*, *Syp*), EC cells (*Chgb, Tph1*, *Tac1* and *Ddc*) and L cells (*Gcg*, *Pyy*, *Insl5*) marker genes were found upregulated (Figure 2C). Of note, neural markers (*Tnr*, *Nrxn1*, *Hap1*, *Lrrn1*, *Ntm*) were observed in Olfr78-expressing cells, pointing out their neuroendocrine identity. GFP^+ve^ cells were especially enriched in presynaptic and postsynaptic markers (*Syt14*, *Snap25*, *Syn1* and *Nlgn2*, *Dlg3*, *Shank2*, respectively) and expressed the neurofilament marker *Nefm* (Figure 2D). These data are coherent with the function of EC cells, reported to establish interactions through neuropod processes with surrounding epithelial cells and synapses with enteric nerve cells (Bellono et al., 2017; Bohórquez et al., 2014, 2015). Moreover, since EEC differentiation involves transient intermediate committed states, each coined by a particular set of transcription factors (Gehart et al., 2019), we investigated the expression of these genes in Olfr78-expressing cells (Figure 2E). GFP^+ve^ cells did not express early EEC markers (*Dll1*, *Isx*) but were enriched in key early/intermediate and intermediate transcriptions factors (TF) (*Pbx1*, *Rybp*, *Sox4* and *Pax4*, *Rcor2*, *Rfx3*, respectively), indicating that Olfr78 transcription initiates in these EEC progenitors. Expression of *Olfr78* was also maintained in intermediate/late and late EEC cells based on enrichment of GFP^+ve^ cells in the following TFs (*Insm1*, *Neurod1*, *Rfx6*, *Lmx1a* as well as *Pax6*, *Egr3* and *Emb*) (Figure 2E and EV2B). Altogether, these data indicated that Olfr78 is expressed in epithelial EEC progenitors during their commitment towards L and EC lineages and in mature EEC subtypes.

**Figure 2.**
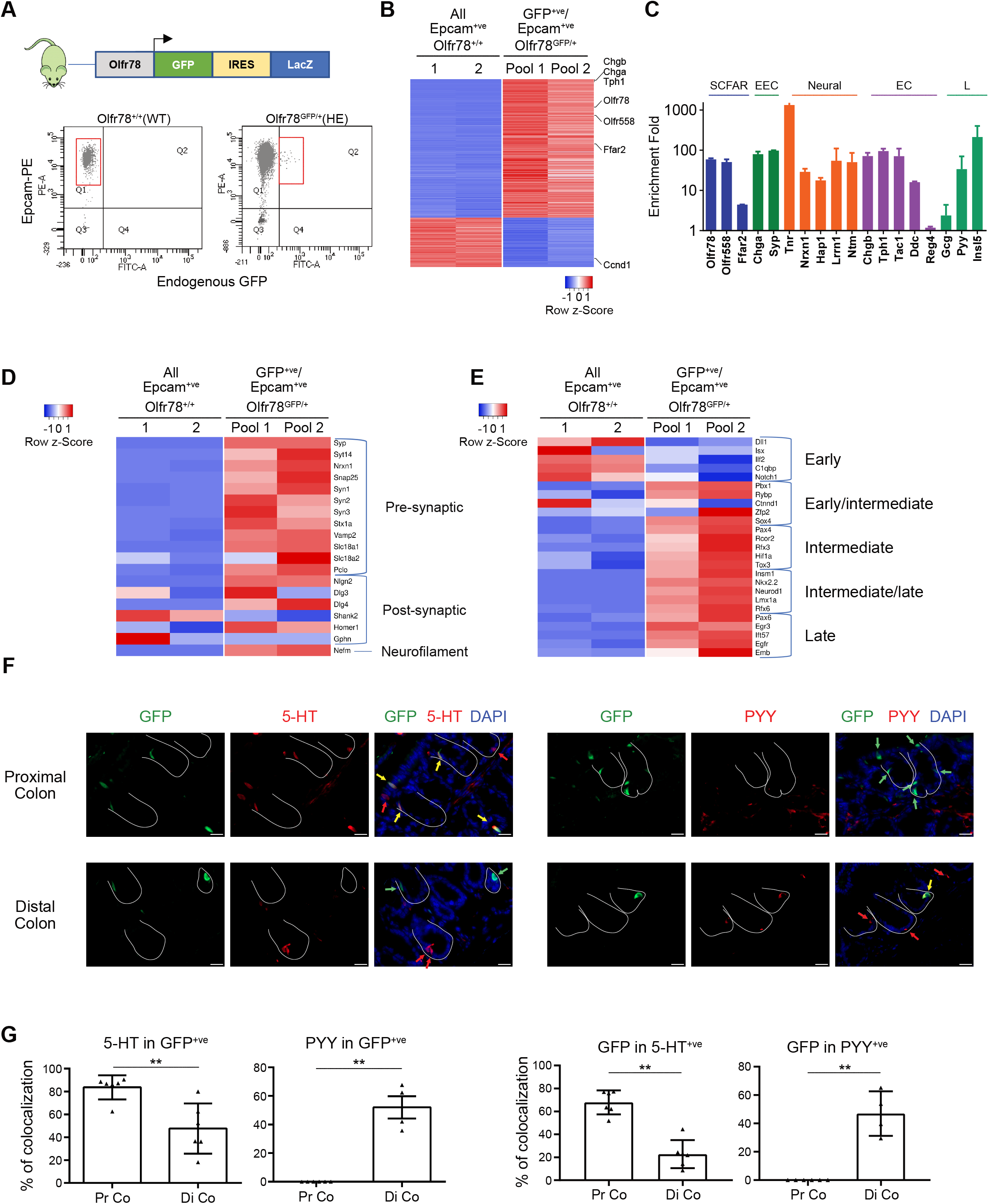
Olfr78 is expressed in different subtypes of enteroendocrine cells in the colon. A. Top: Schematic representation of the Olfr78-GFP knockin/knockout mouse line. Bottom: strategy of isolation and sorting by flow cytometry of Epcam^+ve^/GFP^+ve^ cells. Graphs show 10,000 cells and 90,000 cells in Olfr78^+/+^ (WT) and Olfr78^GFP/+^ (heterozygous, HE) samples, respectively. B. Heatmap of differentially regulated genes in colon Epcam^+ve^/GFP^+ve^ cells (each pool obtained from 2 Olfr78^GFP/+^ mice) versus Epcam^+ve^ cells (2 biological replicates from a single WT mouse) showing log_2_ fold change < −2 or > 2 and false discovery rate ≤ 0.001. C. Histogram showing fold change of expression of relevant marker genes in Epcam^+ve^/GFP^+ve^ cells vs Epcam^+ve^ cells. D. Heatmap of differentially regulated pre-synaptic, synaptic and neuropod-associated genes in Epcam^+ve^/GFP^+ve^ cells vs Epcam^+ve^ cells. E. Heatmap of differentially expressed transcription factors involved in EEC differentiation in Epcam^+ve^/GFP^+ve^ cells vs Epcam^+ve^ cells. F. Immunofluorescence showing 5-HT or PYY production in GFP^+ve^ cells in proximal and distal colon of Olfr78^GFP/+^ mice. Crypts are delineated in white. Red and green arrows point to cells expressing only hormones or GFP, respectively. Yellow arrows indicate GFP^+ve^ cells colocalizing with 5-HT or PYY. Nuclei were counterstained with DAPI. G. Quantification of GFP^+ve^ cells expressing 5-HT or PYY in proximal and distal colon (Pr Co and Di Co, respectively) or 5-HT^+ve^ and PYY^+ve^ cells expressing GFP. Each symbol indicates the value of a given mouse. Data information: Scale bars = 20 µm (F). Data are represented as mean ± SEM. ** P <0.01, Mann Whitney test (G).

We confirmed these results on colon tissues of Olfr78-GFP HE mice by immunofluorescence studies. Indeed, the GFP reporter was detected in the EC and L cell lineages (in accordance with 5-HT and PYY expression, respectively) (Figure 2F, G). Interestingly, GFP was expressed in L (PYY^+ve^) cells exclusively located in the distal colon, in agreement with a previous report (Billing et al., 2019). In contrast, GFP expression in EC (5-HT^+ve^) cells was predominant in the proximal colon (Figure 2F, G). Overall, these results revealed a regional expression pattern of colon epithelial Olfr78-expressing cells in EEC subtypes.

### Loss of Olfr78 impairs terminal differentiation into enterochromaffin cells

To investigate whether Olfr78 could play any role in the colon, we took advantage of the Olfr78-GFP knockin/knockout mouse line. First, we confirmed that Olfr78-GFP homozygous mice were knockouts (KO) in the whole colon by qRT-PCR (using primers targeting the coding exon) and by RNAscope (Figure 3A and EV3A). At the histological level, loss of Olfr78 did not significantly affect Goblet cell differentiation or cell proliferation in colon (Figures 3B and EV3B). After having checked that GFP^+ve^ cells were detectable in the colon of Olfr78-GFP KO (Figure EV3C), we sorted by flow cytometry colon Epcam^+ve^/GFP^+ve^ cells from these mice (2 pools, each obtained from 2 individual mice) (Figure 3C). Then, we compared their transcriptome obtained by RNAseq to that of Epcam^+ve^/GFP^+ve^ cells from Olfr78-GFP HE mice (Figure 3D, upper panel). This resulted in a list of 364 genes up- and 1174 genes downregulated in epithelial GFP KO vs HE cells. Strikingly, Olfr78 deficiency correlated with de-enrichment in biological processes associated with neurogenesis, regulation of secretion and synapse organization (downregulated pre-synaptic and post-synaptic associated genes are listed in Figure EV3D) meanwhile cell division, DNA replication and cellular response to stress were upregulated in GFP^+ve^ KOs vs HEs (Figure 3D, lower panel). Next, focusing on the EEC subtypes previously identified as expressing the Olfr78 receptor, we observed significant downregulation in EC markers involved in serotonin production and metabolism (*Tph1*, *Ddc, Gch1*), granule secretion (*Chgb*, *Chga, Rab3c, Gstt1*), and lineage commitment (*Lmx1a*, *Fev*) in GFP^+ve^ KO cells (Figure 3E). In contrast, similar comparison on L cell marker genes (*Pyy*, *Gcg*, *Insl5*, *Etv1*) did not evidence any clear differential expression pattern in GFP^+ve^ KO cells (Figure 3E). These data suggested that the loss of Olfr78 expression in EEC precursors was specifically interfering with the EC subtype terminal differentiation. Downregulation of *Chgb* expression, but not *Pyy*, was confirmed by qRT-PCR on the whole proximal colon exclusively (Figure 3F). The observation that *Tph1* was not found downregulated on whole tissues might be explained in part by the fact that this gene, coding for the rate-limiting enzyme in serotonin synthesis, is still expressed in about 35% of the EC cells that do not express Olfr78 (Figure 2G). To our surprise, expression of the cognate SCFA receptor Olfr558 dropped down to 19% residual levels in Olfr78-deficient tissues (Figure 3F). Accordingly, the numbers of 5-HT^+ve^ and Olfr558^+ve^ cells in proximal colon were significantly reduced (by 26% and 74%, respectively) in the absence of Olfr78 expression (Figure 3G). To fully decipher the identity of epithelial GFP^+ve^ KO cells, we performed single-cell RNAseq (scRNAseq) on colon cells from Olfr78-GFP WT and Olfr78-GFP KO mice. After data merging using the Seurat package, we isolated and clustered EECs defined by *Chga* expression in epithelial cells (Figure 3H). In the Olfr78 WT sample, as expected, the EEC cluster was constituted of 2 groups, defined as EC and L cells, based on specific marker genes expression (Figure 3I, J). In the Olfr78 KO sample, the L cell group was also present and exhibited a transcriptome similar to that of WT L cells. In contrast, the EC group was extremely reduced in Olfr78-deficient tissues and appeared to be replaced by a third cluster of cells in an “undefined state” (Figure 3I, J). Indeed, this later group of cells expressed low levels of EC marker genes (such as *Tph1*, *Chgb* or *Ddc*) and higher levels of non-EEC genes, such as *Sycn* involved in pancreatic acinar cell exocytosis, *Mfsd4a* an atypical solute carrier transporter expressed in several brain areas and *Tfdp2* a transcription factor involved in cell cycle control (Asle et al., 2005; Chen & Lodish, 2014; Perland et al., 2017). In addition, *Agr2* and *Muc2*, two early goblet cell differentiation markers, were also enriched in Olfr78 KO cells. Altogether, these scRNAseq studies further indicated that loss of Olfr78 in mouse colon results in improper EC differentiation, blocking cells in an undefined state, with characteristics of secretory lineage identity.

**Figure 3.**
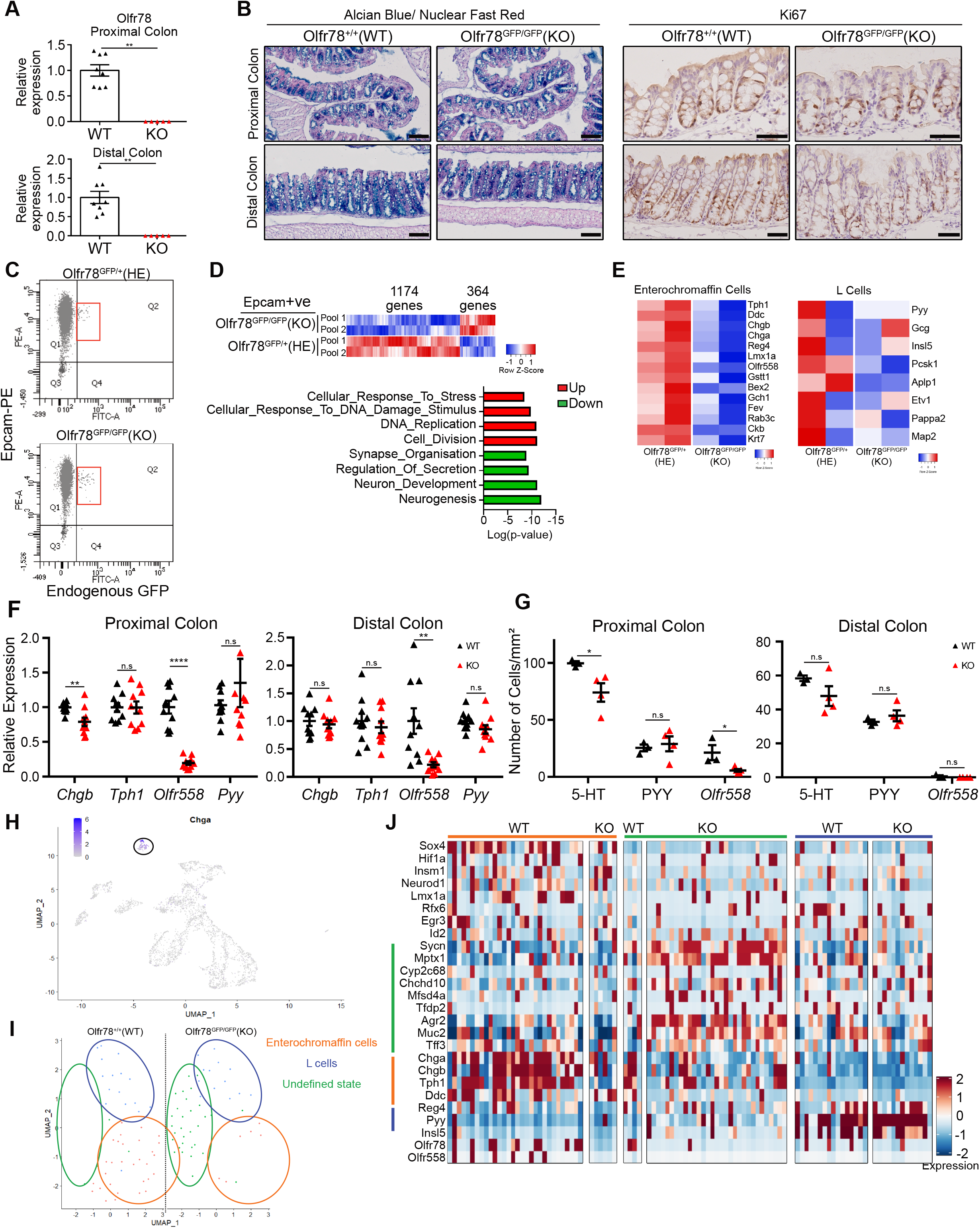
Loss of Olfr78 impairs terminal differentiation into enterochromaffin cells. A. Analysis of Olfr78 expression by qRT-PCR in proximal and distal colon from WT and Olfr78-GFP KO mice. Relative expression levels were arbitrary set to 1 in WT samples. Each symbol indicates the value of a given mouse (n = 8 WT, 5 KO). B. Representative images showing Alcian Blue-Nuclear Fast Red staining to evidence Goblet cell differentiation (left) and immunohistochemistry for Ki67 staining (right) on proximal and distal colon in Olfr78-GFP WT and Olfr78-GFP KO mice. C. Bottom: strategy of isolation and sorting by flow cytometry of Epcam^+ve^/GFP^+ve^ cells. Graphs show 90,000 cells in Olfr78^GFP/+^ (HE) and Olfr78^GFP/GFP^ (homozygous/knockout, KO) samples, respectively. D. Transcriptome comparison between colon epithelial Olfr78-GFP KO vs Olfr78-GFP HE GFP^+ve^ cells. Upper panel: heatmap of differentially expressed genes between Epcam^+ve^/GFP^+ve^ cells coming from 2 pools of 2 different Olfr78-GFP HE or KO mice. The number of genes differentially modulated is indicated. Lower panel: GSEA-Biological processes for upregulated (red) and downregulated (blue) gene lists in Olfr78-GFP KO vs Olfr78-GFP HE GFP^+ve^ cells. E. Heatmaps showing expression levels of EEC markers in Olfr78-GFP HE and Olfr78-GFP KO GFP^+ve^ pools. F. Expression of EEC markers analyzed by qRT-PCR on Olfr78-GFP WT and Olfr78-GFP KO proximal and distal colon biopsies. Relative expression levels were set to 1 in WT samples. Each symbol indicates the value for a given mouse (n = 10 WT, 10 KO). G. Quantification of 5-HT, PYY and *Olfr558* expressing cells in proximal and distal colon of Olfr78- GFP WT and Olfr78-GFP KO mice performed based on IHC staining (5-HT and PYY) or RNAscope (*Olfr558*). Each symbol indicates the value of a given mouse (n = 3 WT, 4 KO). H. UMAP of merged epithelial cells from WT (n= 1,764 cells) and KO (n= 1,500 cells) mice after scRNAseq. The cell cluster showing *Chga* expression is circled. I. UMAPs of WT and KO *Chga*-expressing EEC cells showing clustering into 3 cell populations. J. Heatmap showing the expression of various genes in the 3 EEC-associated groups identified in I.. Data information: Scale bars = 100 µm (Alcian Blue) or 50 µm (Ki67) (B). Data are represented as mean ± SEM. (A) Mann-Whitney tests: ** P=0.0016 (Pr Co and Di Co). (F) Unpaired t-tests (Tph1 in Pr Co; Chgb, Tph1 and Pyy in Di Co), unpaired t-tests with Welch’s correction (Chgb,and Olfr558 in Pr Co and Olfr558 in Di Co), Mann-Whitney test (Pyy in Pr Co). n.s = not significant; ** P <0.01; *** P <0.001 (G) Unpaired t-tests (5-HT and PYY in Pr Co), Mann-Whitney test (Olfr558 in Pr Co and 5-HT, PYY and Olfr558 in Di Co). n.s = not significant; * P <0.05.

### Terminal differentiation into serotonin-producing cells is regulated by epithelial Olfr78 expression

Since Olfr78-expressing cells were detected in epithelium and mesenchyme throughout the colon, we sought to further investigate if the phenotype observed in full Olfr78-GFP KO mice was related to loss of expression in the epithelium, mesenchyme or both compartments. For this purpose, we generated a new mouse line harboring an Olfr78 floxed allele targeting the coding exon, thereafter referred to as Olfr78^fx^ (Figure 4A). To study the impact of epithelial ablation of Olfr78 on EEC differentiation, Olfr78^fx^ mice were crossed with Vil1^Cre/+^ mice (where Cre recombinase is expressed under the control of the Villin promoter, active in epithelial cells) to generate Vil1^Cre/+^-Olfr78^fx/fx^ mice, referred to as Olfr78 eKO (eKO, for KO in epithelium). First, we validated the efficient deletion of the targeted region on DNA isolated from colon biopsies of Vil1^Cre/+^-Olfr78^fx/fx^ (Figure EV4A). We also confirmed by RNAscope that loss of Olfr78 was restricted to the epithelium, with non-epithelial expression being preserved and representing 12% residual *Olfr78* expression in proximal colon by qRT-PCR analysis (Figure 4B, C). Then, we investigated colon histology and mucus production by Alcian Blue staining and did not notice any significant altered Goblet cell differentiation or change in epithelial cell proliferation (Figures 4D and EV4B). Next, EEC cell differentiation was studied by qRT-PCR using EC and L cell marker genes. As observed in Olfr78-GFP KO mice, Olfr78 eKO mice exhibited a consistent reduction in the EC markers *Chgb* and *Olfr558* (by 35% and 48%, respectively) in proximal, but not distal, colon (Figure 4E). Instead, expression levels of the L cell marker *Pyy* were not different between the genotypes (Figure 4E). Moreover, in agreement with qRT-PCR analyses, the density in EC cells was decreased by 32% in the proximal colon of eKO mice as compared to littermate controls whereas no change in EC and L cells density was observed in distal colon (Figure 4F). Altogether, these findings further demonstrated that epithelial expression of Olfr78 in EECs is necessary to generate mature EC cells in proximal colon. To determine whether Olfr78 loss could alter colon 5-HT levels, we performed dosage of this hormone in stools collected from controls and eKO mice. Consistent with the overall decrease in EC density observed in eKO mice, 5-HT concentration was found tendentially reduced by 25% in the absence of this SCFA receptor (p-value = 0.0775) (Figure 4G). Finally, to investigate the molecular mechanisms by which Olfr78 can contribute to EC differentiation, we used a 3D organoid culture model (Sato et al., 2011). Following 7 days of culture after replating, Olfr78-GFP WT or Olfr78- GFP KO colon organoid lines (generated from individual mice) were stimulated for 48 hours with acetate at a concentration of 20 mM. As shown in Figure 4H, expression of *Chgb*, used as one of the most specific markers of EC maturation, but not the L cell marker *Pyy*, was upregulated by 40% in WT organoids upon acetate challenge, showing that this SCFA stimulates EC differentiation. In contrast, expression of *Chgb* was downregulated by 20% in Olfr78-GFP KOs following acetate challenge as compared to untreated conditions (Figure 4H). Together, these data indicated that maintenance of EC maturation involves activation of the epithelial Olfr78 receptor via its ligand acetate in the mouse colon.

**Figure 4.**
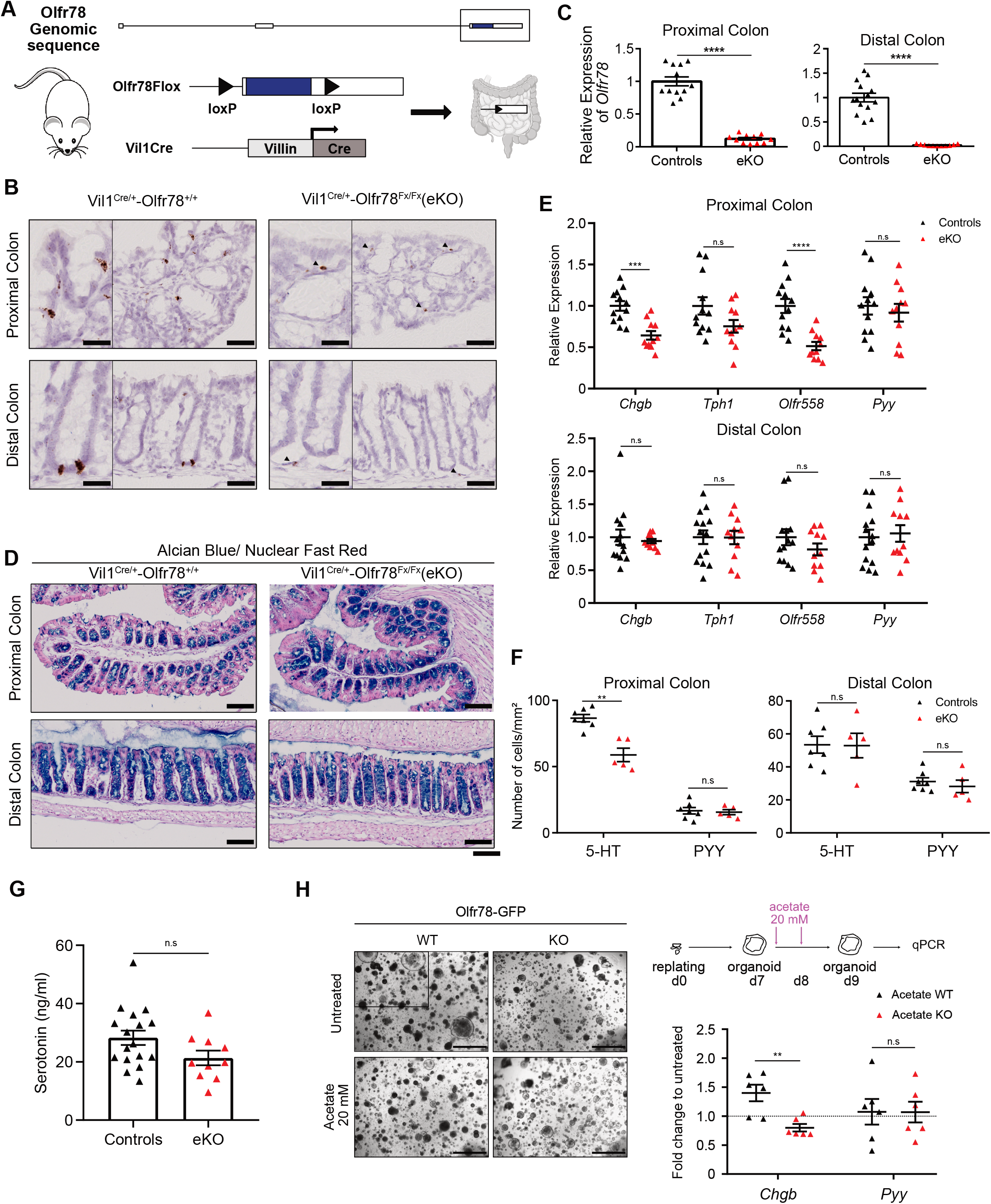
Terminal differentiation into serotonin-producing cells is regulated by epithelial Olfr78 expression. A. Genomic construction of the Olfr78^fx^ line. The LoxP sites flanking the exon 3 coding for Olfr78 are evidenced. The coding sequence is labeled in blue. B. Representative pictures of RNAscope staining showing specific epithelial ablation of *Olfr78* expression in Vil1^Cre/+^-Olfr78^fx/fx^ (eKO) mice but not in Vil1^Cre/+^-Olfr78^+/+^ mice. Arrowheads show non-epithelial expression of *Olfr78*. C. Analysis of residual *Olfr78* expression by qRT-PCR in control or eKO colon biopsies. Relative expression levels were arbitrary set to 1 in control samples. Each symbol indicates the value of a given mouse (n= 12 controls and 11 eKO). Controls corresponded to Vil1^+/+^-Olfr78^fx/fx^ and Vil1^Cre/+^-Olfr78^+/+^ mice. D. Representative pictures of Alcian Blue-Nuclear Fast Red staining on proximal and distal colon from Vil1^Cre/+^-Olfr78^+/+^ or Vil1^Cre/+^-Olfr78^fx/fx^. E. Expression of EEC markers was analyzed by qRT-PCR on colon biopsies from controls or eKO mice. Relative expression levels were arbitrary set to 1 in controls. Each symbol indicates the value for a given mouse (n= 12-14 controls, 11 eKO). F. Quantification of 5-HT and PYY-expressing cells in proximal and distal colon tissues from controls or eKO mice performed based on IHC staining. Each symbol indicates the value of a given mouse (n = 7 controls, 5 eKO). G. Serotonin dosage in stools collected from controls and eKO colon (mice age: between 8 and 27 weeks-old). Each symbol indicates the value of a given mouse (n = 17 controls, 10 eKO). H. Left panel: Representative pictures of Olfr78-GFP WT and Olfr78-GFP KO colon organoid cultures after 48hours of treatment (acetate 20 mM) or in untreated conditions. Right panel: expression levels of EEC markers analyzed by qRT-PCR on colon organoids after 48 hours of treatment. Data are reported as the fold-change of expression in treated vs control conditions in paired samples. Each symbol indicates the mean value of three independent experiments performed on organoids, each line being generated from a given mouse (n= 6 WT and 6 KO). Data information: Scale bars: 50 µm (B, large view) or 25 µm (B, inset), 100 µm (D), 1 mm (H). Data are represented as mean ± SEM. (C) Unpaired t-tests with Welch’s correction, **** P <0.0001. (E) Unpaired t-tests (Pr Co), Mann-Whitney test (Di Co). n.s = not significant; *** P <0.001; **** P <0.0001. (F) Mann-Whitney tests, n.s = not significant, ** P <0.01. (G) Unpaired t-test (p=0.075). (H) Mann-Whitney test, n.s = not significant, ** P <0.01.

### Loss of Olfr78 expression alters colon homeostasis

Having provided evidence that epithelial Olfr78 regulates the production of functional EC cells known to contribute to colon physiology, we investigated whether absence of Olfr78 could affect global colon homeostasis. First, we analyzed the whole epithelium by studying the transcriptome of isolated crypts from Olfr78-GFP WT and Olfr78-GFP KO mice (n = 2 and 5, respectively) by bulk RNA seq. Analysis of differentially expressed genes with a FDR ≤ 0.01 resulted in a list of 261 up- and 560 downregulated genes in Olfr78-GFP KO vs WT crypts (Figure 5A). MDS plots showed separate clustering of both genotypes (Figure 5A). *In silico* studies on modulated biological processes and C8 cell type signature gene sets (GSEA) revealed upregulation of genes involved in cytoskeleton organization, intracellular transport, or cell morphogenesis processes. Moreover, genes associated with aerobic cellular respiration, response to oxidative stress or monocarboxylic acid metabolic processes were downregulated (Figure 5B). Indeed, genes involved in mitochondrial function and genes belonging to several antioxidant families (such as peroxiredoxins, glutathione-S-transferases and glutathione peroxidases) as well as *Nfe2l2*, a major transcription factor regulating expression of some of these genes, were found downregulated in Olfr78-GFP KO crypts as compared to controls (Figure 5C). In contrast, expression of *Duox2*, which allows production of the oxidative molecule H_2_O_2_, was increased in Olfr78-GFP KO crypts (Figure 5C). RNAseq data also suggested that absence of Olfr78 expression had altered cell fate in the epithelium, with an increased proportion of cells being engaged into the neuroendocrine fate at the expense of the other cell types (Figure EV5A). The significant increase in density of GFP^+ve^ cells observed in Olfr78-GFP KO vs Olfr78-GFP HE colons, secretory in their identity, further sustained this idea (Figure 5D). Regarding the surrounding non-epithelial cells present in colon, analysis of scRNAseq data from Olfr78-GFP WT and Olfr78-GFP KO mice indicated that they were clustered into 11 distinct populations in both genotypes, without major quantitative differences detected in immune CD45^+ve^ cells or stromal Pdgfra^+ve^ cells in the absence of Olfr78 (Figure EV5B-D). Next, since 5-HT production and its luminal release is associated with microbiome homeostasis, we studied the impact of Olfr78 loss on fecal microbiota by performing a metagenome analysis on colon stools from 7 Olfr78-GFP WT and 4 Olfr78-GFP KO mice (Figure 5E). Interestingly, the *Firmicutes*/*Bacteroidetes* ratio, proposed as marker of dysbiosis when dysregulated, was significantly increased in Olfr78-GFP KO vs WT samples, despite no significant difference in total body weight between genotypes (Figures 5F and EV5E). We also noticed the virtual absence of *Turicibacter sanguinis* in Olfr78-GFP KO samples (Figures 5G and EV5F), a species reported to take advantage of luminal 5-HT released by ECs to increase its fitness and colonization ability in the colon (Fung et al., 2019). Knowing that microbiota are the main producers of SCFAs in the colon and having found that Olfr78-deficient mice had signs of dysbiosis, we analyzed the concentration of various SCFA compounds (from C2 to C6) in the stools of Olfr78-GFP WT and KO mice. No significant differences were detected between genotypes for any of the SCFAs analyzed, including acetate and propionate, the two reported ligands of Olfr78 (Figure EV5G). In summary, our results indicate that the loss of Olfr78 receptor alters colon homeostasis, characterized by deficient epithelial detoxification potential and mild dysbiosis under chow diet.

**Figure 5:**
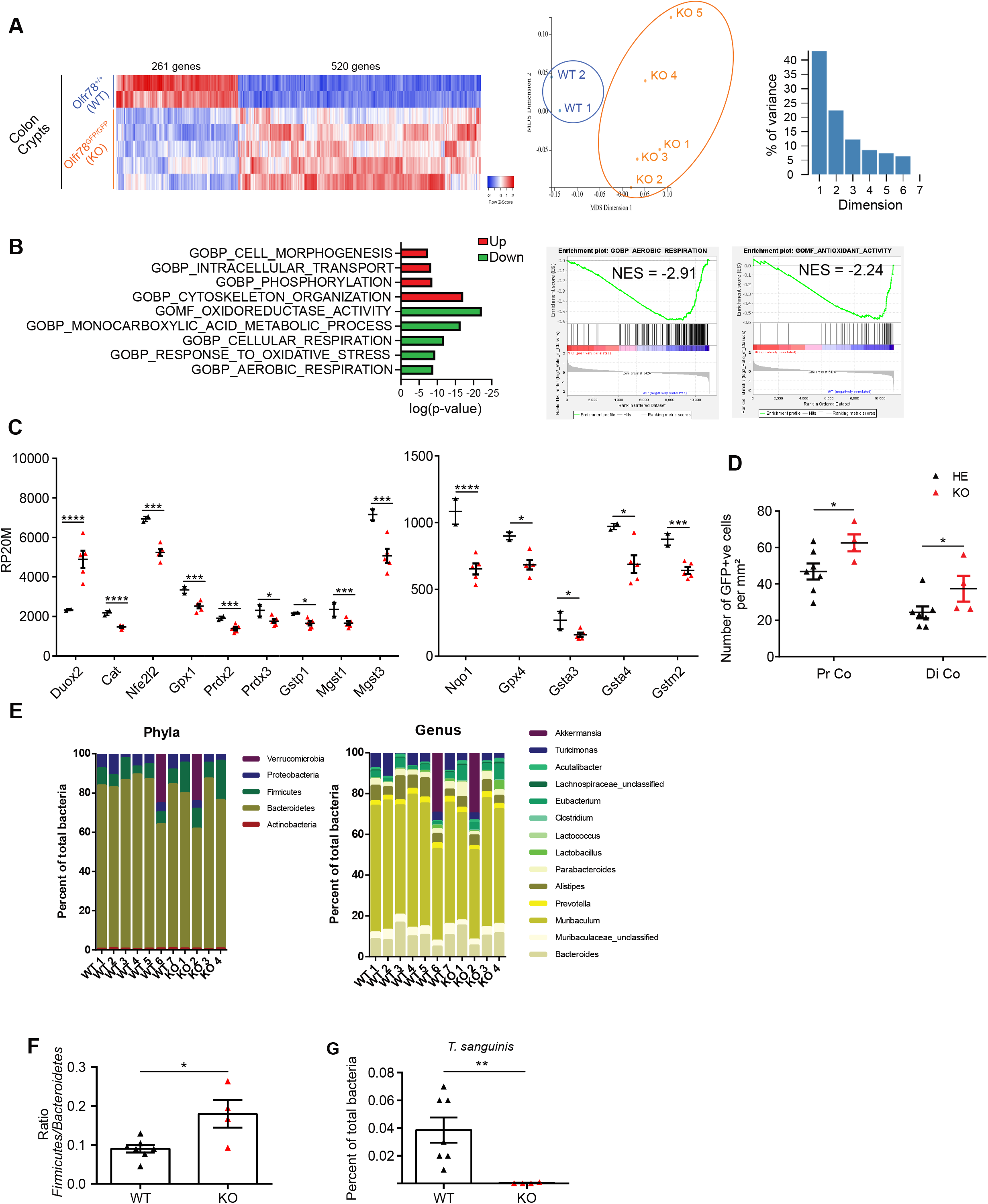
Loss of Olfr78 expression alters colon homeostasis. A. Left: Heatmap of differentially expressed genes identified by RNAseq on colon crypts isolated from Olfr78-GFP WT and KO mice (n = 2 and 5, respectively) (FDR ≤ 0.01 from Degust). Right: Multi-dimensional scaling (MDS) plot of RNA-seq datasets on genes expressed at least on 2 samples and minimum 40 reads per 20 million reads. B. Left: Modulated Mol-Sig GSEA Biological processes in the transcriptome of Olfr78-GFP KO vs Olfr78-GFP WT crypts. Right: GSEA showing de-enrichment of the Antioxidant activity and Aerobic respiration datasets in Olfr78-GFP KO vs Olfr78-GFP WT crypts. NES: Normalized Enrichment Score. C. Graphs showing expression levels in RP20M (Reads per 20 million mapped reads) of some target genes associated with the pathways identified in B. D. Quantification of GFP^+ve^ cell density in proximal (Pr Co) and distal colon (Di Co) from Olfr78-GFP HE and Olfr78-GFP KO mice performed based on IHC staining. Each symbol indicates the value of a given mouse (n = 7 HE, 4 KO). E. Histograms showing the relative microbial abundance at the Phyla and Genus levels in Olfr78-GFP mice (n = 7 WT and 4 KO) by metagenome sequencing. F. Ratio of the prevalence of *Firmicutes*/*Bacteroidetes* populations obtained from the data in E. Each symbol indicates the value of a given mouse (n = 7 WT, 4 KO). G. Percentage of prevalence of *Turicibacter sanguinis* obtained from the data in E. Each symbol indicates the value of a given mouse (n = 7 WT, 4 KO). Data information: Data are represented as mean ± SEM. (C) * FDR <0.05; *** FDR <0.001; **** FDR <0.0001, EdgeR method from Degust. (D, F, G) Mann-Whitney tests, n.s = not significant; *P <0.05; ** P <0.01.

## DISCUSSION

Since their discovery in the olfactory epithelium in 1991 (Buck & Axel, 1991), several olfactory receptors have been reported to be ectopically expressed (Lee et al., 2019). Among these receptors, Olfr78/OR51E2 expression was detected in the gut at significant levels (Billing et al., 2019; Fleischer et al., 2015; Kotlo et al., 2020; Lund et al., 2018). In the present study, we have investigated the biological function of Olfr78 in the colon and revealed that this receptor regulates enterochromaffin cell maturation under homeostatic conditions.

In line with previous studies, using the reporter Olfr78-GFP mouse line, transcriptomic and immunostaining methods, our work confirmed that Olfr78 is mainly present in two epithelial enteroendocrine cell subtypes in colon: EC and L cells, characterized by the expression of the marker genes *Tph1*, *Ddc*, *Tac1* and *Pyy*, *Insl5*, respectively (Billing et al., 2019; Lund et al., 2018). Olfr78 expression was detected in the majority in EC cells (80%) of the proximal colon and was identified in both EC and L cells in the distal colon. By using the resource generated by Gehart et al (2019) on time-resolved single-cell transcriptional mapping of enteroendocrine differentiation in the small intestine, we further determined by bulk and single cell RNA seq methods that expression of Olfr78 starts in early/intermediate EEC precursors (*Rybp*, *Sox4*) and is maintained throughout differentiation in intermediate/late (*Lmx1a*, *Rfx6*) and late (*Pax6*) stages. Analysis of Olfr78-deficient cells revealed that the absence of this receptor leads to markedly reduced EC cell differentiation, characterized by downregulation of the *Lmx1a* and *Fev* transcription factors (Wang et al., 2010) while no significant decrease in the expression of the transcription factor *Rfx6*, promoting differentiation into L cells, was observed (Piccand et al., 2019). More globally, dysregulated EC differentiation in Olfr78-deficient cells leads to downregulation of neuronal genes, as represented by genes encoding pre-synaptic and secretion granule components. In this regard, the human ortholog OR51E2, sharing 93% identity with Olfr78, was recently detected in human pulmonary neuroendocrine cells and reported to activate a neuroendocrine phenotype in prostate cancer cells (Abaffy et al., 2018; Kuo et al., 2022). Taken together with our results, these data suggest that, in response to its ligands, the Olfr78/OR51E2 receptor can promote a neuronal-like phenotype outside the olfactory epithelium. Interestingly, a study recently revealed that SCFA ligands (acetate or propionate), used at concentrations like those found in the colon (10mM), promote EC cell differentiation in colon organoid cultures, although the identity of the transducing receptor(s) was not investigated (Tsuruta et al., 2016). In the present work, we provide further evidence that EC cell maturation involves, at least in part, the Olfr78 receptor, through the activation by acetate, one of its ligands. Regarding the L cell subtype, their density and the expression levels of the anorexigenic PYY peptide in the colon did not appear modified in absence of Olfr78 expression. However, it is not excluded that this receptor could regulate secretion of this hormone in L cells as proposed by Nishida et al. (2021)

Despite substantial altered EC identity detected in Olfr78-deficient cells in both transgenic lines, this resulted in a consistent but modest reduction (by 30%) in the number of serotonin-producing EC cells in the proximal colon and a 25% reduction in fecal 5-HT levels. Moreover, expression of the *Tph1* gene, coding for the rate-limiting enzyme in 5-HT synthesis, did not appear downregulated at the whole tissue level. Several hypotheses, not mutually exclusive, could be proposed to explain these observations. Firstly, 35% of EC cells do not express Olfr78 in the proximal colon, and can *a priori* differentiate normally, being able to produce some 5-HT. Secondly, in the small intestine, Haber et al. (2017) reported some expression of *Tph1* in early EEC precursors. In colon EEC precursors, a similar event may contribute to dampen the observed reduction of *Tph1* expression in Olfr78-deficient EC cells. The major impact of Olfr78 loss detected at the whole tissue level was on chromogranin’s expression, especially chromogranin B. This later is a component of secretory granules recently reported to form a chloride channel involved in the secretion of neurotransmitters (Yadav et al., 2018). It is expected that the reduction in chromogranin B levels in Olfr78-deficient cells would also alter the 5-HT secretion process.

The present study also sheds light on the complex expression profile of SCFA receptors in colon where the concentration of their ligands is the highest. Indeed, Olfr558 and Ffar2 receptors were found enriched in Olfr78-expressing cells. This suggests different genetic and/or epigenetic regulatory mechanisms of olfactory receptors’ expression in EC as compared to olfactory sensory neurons where one neuron expresses only one receptor (Serizawa et al., 2004). Moreover, we unexpectedly found that the loss of Olfr78 induced a decrease in Olfr558 expression in EC cells. Although not formally excluded, the likelihood that this would result from an artifact of the transgenic strategy, affecting the genetic locus that bears the two genes, remains low since the effect was observed in both mouse lines. Regarding the hypothetical function of Olfr558 in colon EC cells, one possibility could be that this receptor exhibits in colon, similar functions as those described in the small intestine (Bellono et al., 2017). Stimulation of Olfr558 with the butyrate and isovalerate ligands also present in colon would promote 5-HT basolateral release from EC cells to activate neuronal circuitries. Further experiments will be needed to investigate this hypothesis. Besides, our study also revealed that Olfr78-expressing EC cells express the Ffar2 receptor. Contrary to Olfr78 and Olfr558, this receptor was also detected in other cells in the bottom of colon crypts, suggesting that it would exert a more general function in epithelial cells than odorant receptors. Previous studies have demonstrated that Ffar2 regulates inflammation in colon (Kim et al., 2013). However, the potential role of Ffar2 in colon EC cells remains to be determined; especially considering that this receptor, like Olfr78, recognizes acetate and propionate as ligands, but that these two receptors induce signaling through opposite cascades involving Gq/Gi and Gs proteins for Ffar2 and Olfr78, respectively (Brown et al., 1989; Saito et al., 2009). Interestingly, it was previously demonstrated that Olfr78 and Ffar3 coordinate their activity to regulate blood pressure (Pluznick et al., 2013), raising the possibility of a similar mechanism involving Olfr78 and Ffar2 in EC cells. Additional studies are needed to explore this hypothesis and to further decipher the complex interplay between both receptors in the regulation of EC cell maturation.

At the global tissue level, we detected that Olfr78-deficient mice had a modified microbiota, an observation also made by Kotlo et al. (2020). Indeed, Olfr78 KO mice exhibit an increased *Firmicutes*/*Bacteroidetes* ratio as compared to WT littermates. In addition, despite the modest reduction in luminal 5-HT levels in Olfr78-deficient mice, *Turicibacter sanguinis*, a bacterium reported to need serotonin to increase its fitness, was virtually absent in Olfr78-GFP KO stools, showing that alteration in the epithelium significantly impacted the microbiota (Fung et al., 2019). Since the *Firmicutes*/*Bacteroidetes* ratio is increased in obesity, it would be interesting to further investigate the impact of Olfr78 loss on the global metabolic status under high fat diet conditions. Finally, our transcriptomic data on colon crypts showed that loss of Olfr78 expression impairs colon epithelium homeostasis by reducing the expression of antioxidant genes as well as genes linked to mitochondrial activity, but without any major impact on the non-epithelial compartments. Of interest, in inflammatory conditions experimentally induced by dextran sodium sulfate treatment, complete absence of Olfr78 expression increases inflammation and impairs efficient tissue regeneration (Kotlo et al., 2020). Knowing that Olfr78 is also expressed in mesenchymal cells in colon, it will be worth addressing the putative role of these cells in tissue homeostasis during regeneration.

In summary, in the present work, we provide evidence that the Olfr78 receptor is expressed in colon enteroendocrine precursors and is required for proper maturation into the EC cell subtype, devoted to serotonin release. Loss of Olfr78 leads to altered 5-HT levels, dysbiosis and modifies the response to oxidative stress in colon crypts. Given that 5-HT represents a major potential pharmacological target in metabolic disorders and inflammatory bowel diseases, further exploration of the role of Olfr78 in epithelial and stromal compartments under pathological conditions will help to fully elucidate the complex mechanisms regulating SCFA receptors, 5-HT secretion, and colon homeostasis.

## MATERIAL AND METHODS

### Experimental animals

Animal procedures complied with the guidelines of the European Union and were approved by the local ethics committee (CEBEA from the faculty of Medicine, ULB) under the accepted protocol 720N. Mice were bred and maintained under a standard 12 hours-light-dark cycle, with water and rodent chow *ad libitum*. Mice strains were B6;129P2-Or51e2tm1Mom/MomJ, (referred as Olfr78-GFP) and B6.Cg-Tg(Vil1-cre)997Gum/J (Vil1Cre) obtained from The Jackson Laboratory, and B6-Olfr78Tm1Mig (referred as Olfr78Fx) generated by Applied StemCell. Primer used for mouse genotyping are listed in Table 1.

**Table 1.**
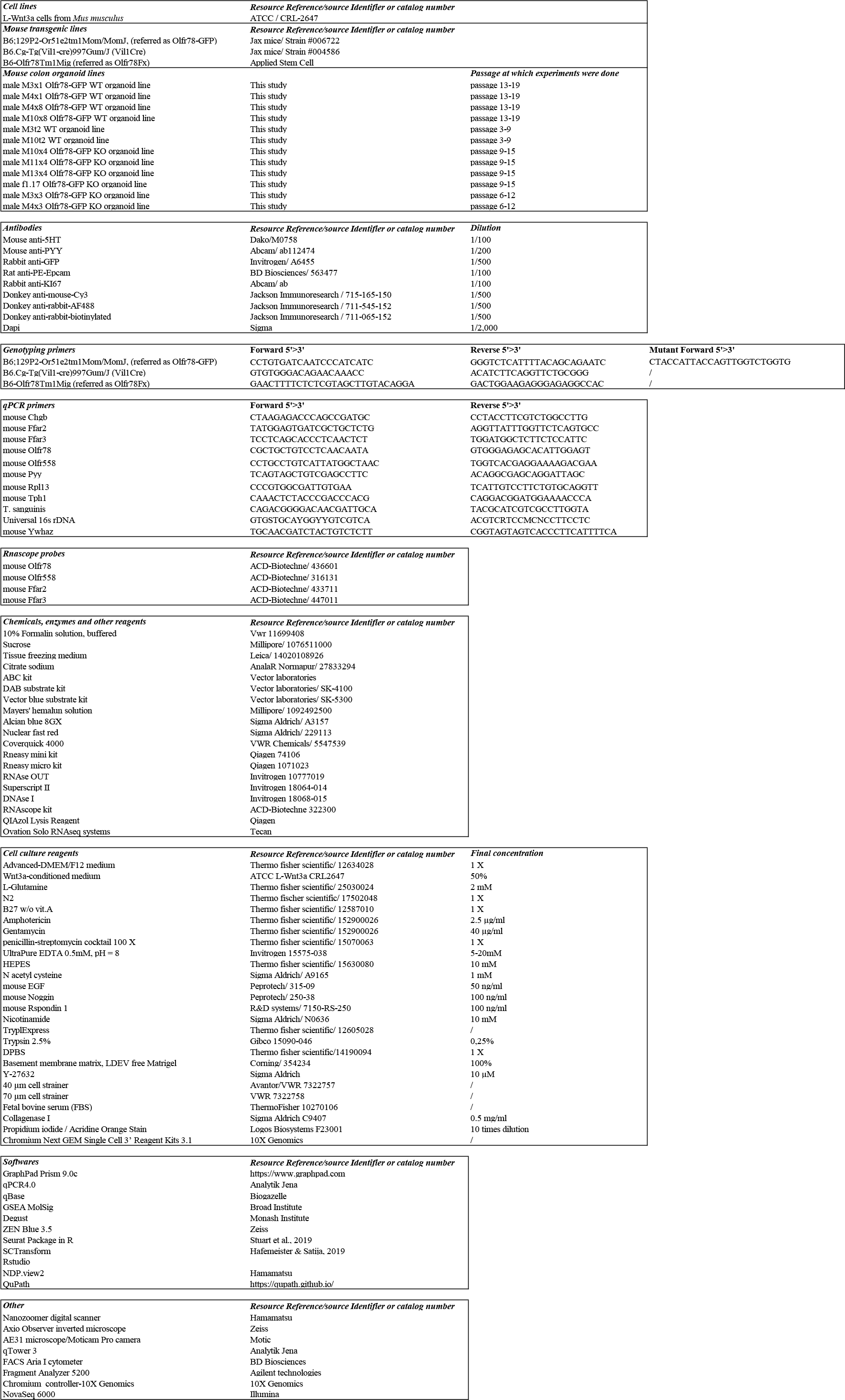

### Tissue processing and immunohistochemical analysis

Intestine samples were immediately fixed with 10% formalin solution, neutral buffered (Sigma-Aldrich) overnight at +4°C and then sedimented through 20% and 30% sucrose solution sequentially (minimum 24 hours each) before OCT (Leica) embedding. Histological and staining protocols as well as immuno-fluorescence/histochemistry experiments were performed on 6 µm sections. Sodium citrate 10 mM, pH 6 was used as epitope retrieval solution. The primary antibodies were incubated overnight at 4°C. The secondary anti-species biotin- or fluorochrome-coupled antibodies were incubated 1 hour at room temperature. The ABC kits (Vector Labs) and substrate Kits (Vector Labs) were used for immunohistochemistry revelation. DAPI or hematoxylin were used for nuclei staining. The primary and secondary antibodies used for staining are listed in Table 1. For the Alcian Blue/Nuclear fast red staining, OCT sections were dried for 20 minutes at room temperature and washed 2 times in PBS for 5 minutes. Slides were incubated for 3 minutes in 3% acetic acid and then for 20 minutes in Alcian blue solution (Sigma Aldrich) at room temperature. Slides were rinsed in 3% acetic acid, running tap water for 2 minutes, followed by distilled water. They were then incubated for 3 minutes in Nuclear Fast Red (Sigma Aldrich) at room temperature and rinsed in running tap water. Slides were mounted in a xylene-based medium (Coverquick 4000, VWR Chemicals) after dehydration or in FluorSave Reagent (Merck). Nanozoomer digital scanner (Hamamatsu) and Zeiss Axio Observer inverted microscope (immunofluorescence) were used for image acquisition. Quantification of epithelial SCFA receptors positive cells along the gut were performed on a minimum of 130 crypt/villus units per sample. Their localization within the colon crypts were quantified on a minimum of 30 cells for each receptor. Colocalization of GFP with 5HT or PYY colon cells was determined on a mean of 50 ± 20 cells per sample and quantification of Ki67^+ve^ cells per crypt was performed on a mean of 27 ± 5 crypts per sample. Cell density for 5-HT/PYY/*Olfr558* or GFP+ve cells per mm^2^ was analyzed on a mean surface of 1.17 ± 0.15 mm^2^ of epithelium delimited by hand on NDP viewer. The number of biological samples (animals) used for each experiment is reported in Figures and Figure legends.

### Crypt isolation and ex vivo culture

The colon was cut in small pieces and put in 20 mM EDTA (Invitrogen) in DPBS (Gibco) for 30 min on ice and shaked at 80 rpm for crypts dissociation. A mechanical dissociation was then performed by ups-and-downs in a FBS pre-coated 10 ml pipette. The mix was passed through a 70 µm filter (Corning) and centrifuged at 300 g for 5 min. At this step, purified crypts were either collected for organoid culture or RNA seq. The organoid culture was performed as described (Sato et al., 2011). Briefly, the medium used consisted in Advanced-DMEM/F12 medium (Gibco) supplemented with 1X GlutaMAX (Gibco), N2 (Gibco) and B27 w/o vit.A (Gibco), gentamycin, penicillin-streptomycin cocktail, amphotericin, 10 mM HEPES (all from Thermofisher scientific), 1 mM N-acetyl cysteine (Sigma Aldrich), 50 ng/ml EGF, 100 ng/ml Rspondin 1 (R&D systems), 100 ng/ml noggin (both from Peprotech), and 10 mM nicotinamide (Sigma Aldrich) and 50% Wnt3a conditioned medium produced by L Wnt-3A cells (ATCC CRL-2647) following manufacturer’s instructions. Culture medium was changed every other day and after 8-9 days in culture, organoids were harvested and digested with TryplExpress (Thermo Fisher Scientific) for 5 min at 37°C. Cells were further mechanically dissociated by ups-and-downs using a 200 µl tips and PBS was added (5-times TrypleExpress volume). Cells were centrifuged at 1,300 rpm for 5 min were (re)plated in Basement membrane matrix, LDEV free Matrigel (Corning). Culture media were supplemented with 10-µM Y-27632 (Sigma Aldrich) in all initial seeding and replating experiments, for 48 hours. Pictures were acquired with a Moticam Pro camera connected to Motic AE31 microscope.

### Gene expression analysis by qPCR and RNAscope

qRT-PCR was performed on total RNA extracted from adult mouse tissues or organoid cultures using the RNeasy Mini Kit (Qiagen). A DNAse I treatment (Invitrogen) was used to remove potential contaminant DNA. cDNA was prepared using RnaseOUT and Superscript II according to the manufacturer’s protocol (Invitrogen). qPCRs were performed on the qTower 3 from Analytik Jena. Gene expression levels were normalized to that of reference genes (Rpl13, Ywhaz) and quantified using the qBase Software (Biogazelle). Primer sequences are reported in Table 1. *In situ* hybridization experiments were performed according to manufacturer instructions with the RNAscope kit (ACD- Biotechne) (probes listed in Table 1).

### Fluorescence-activated cell sorting (FACS) of Olfr78-GFP^+ve^ cells

Colon crypts from 4 to 7 months old mice were isolated as described above and then resuspended in 4 ml of TryplExpress (ThermoFisher Scientific) for 15 min at 37°C under agitation at 75 rpm. Ups-and-downs were performed with a P1000 pipette and incubated for a further 25 min at 37°C and 75 rpm. After a second round of ups-and-downs, 8 ml of binding buffer [PBS-2 mM EDTA-2% BSA (Sigma Aldrich) (w/v)] were added and the mix was passed through a 40 µm filter (Avantor). Cells were centrifuged at 1,300 rpm for 10 min and the pellet was washed with 4 ml of binding buffer. After centrifugation at 1,300 rpm for 3 min, cells were incubated with an Phycoerythrin-coupled anti-Epcam antibody (BD Biosciences) at 1/100 (v:v) in binding buffer for 30 min on ice. After 3 washes with binding buffer, cells were sorted using a FACS Aria I cytometer (BD Biosciences). FSC and SSC intensities were used for debris and doublets’ exclusion. GFP^+ve^ cells were identified using the FITC channel (Olfr78-GFP line) and Epcam through the PE channel. Epcam^+ve^/GFP^+ve^ cells were sorted from 2 pools of 2 HE (1028 and 2511 cells for each pool) and 2 KO mice (1123 and 1506 cells for each pool). Five thousand Epcam^+ve^/GFP^-ve^ cells were collected from a WT mouse as 2 individual samples. The cells were collected in QIAzol lysis reagent (Qiagen).

### Bulk RNA sequencing and Gene Set Enrichment Analysis (GSEA)

RNA was extracted using Rneasy mini kit or miRNeasy microkit (Qiagen) for crypts and sorted cells, respectively, following manufacturer’s instructions, including the on-column DNAse step. RNA quality was checked using a Fragment Analyzer 5200 (Agilent technologies). Indexed cDNA libraries were obtained using the Ovation Solo RNAseq systems (Tecan) following manufacturer recommendations. The multiplexed libraries were loaded on a NovaSeq 6000 (Illumina) and sequences were produced using a 200 Cycles Kit. Paired-end reads were mapped against the mouse reference genome GRCm38 using STAR software to generate read alignments for each sample. The annotation files Mus_musculus.GRCm38.90.gtf was obtained from ftp.Ensembl.org. After transcripts assembly, gene level counts were obtained using HTSeq. For the RNAseq performed on purified colon crypts, differentially expressed genes with minimum 2 CPM (count per million) in minimum 2 samples were identified with EdgeR method (FDR < 0.01) and further analyzed using GSEA MolSig (Broad Institute) (Subramanian et al., 2005). For the RNAseq performed on FACS-sorted cells (Figure 2), differentially expressed genes with minimum 2 CPM in minimum 2 samples and log_2_ fold change > 2 or > −2 were identified with EdgeR method (FDR < 0.001). For the RNAseq performed on FACS-sorted cells (Figure 3), differentially expressed genes with more than 2 CPM in all the samples were analyzed as (KO mean - WT mean)/[(KO SD + WT SD)/2] is < −2 or > 2 and log2-fold change > 0.585 or < −0.585. GSEA analysis on these samples was performed using the identified DEGs.

### Single cell RNAseq (scRNAseq)

Colon tissues from one wild type and one Olfr78-GFP KO mice (respectively 8 and 6 months-old) were cut in small pieces and digested with 0.5 mg/ml Collagenase I (Sigma Aldrich) for 25 min at 37°C under agitation at 75 rpm. Ups-and-downs were performed with a 10 ml pipette and the samples were incubated for a further 20 min at 37°C and 75 rpm. An equal volume of PBS-5 mM EDTA was added, and cells were centrifuged for 10 min at 50 g. Pelleted cells were digested with 5 ml of Trypsin 0.25% (Gibco) for 25 min at 37°C and 75 rpm. Cells were pipetted up and down, passed through a 40 µm filter and pelleted at 50 g for 10 min. The pellet was washed 3 times with PBS-BSA 0.04% (w/v). Cell viability was evaluated by propidium iodide/Acridine Orange staining (85.9% and 90.3% viability in the WT and the KO, respectively) before processing through the Chromium Next GEM Single Cell 3’ Reagent Kits V3.1 (10X Genomics) and sequenced on a Novaseq 6000 (Illumina).

Data were analyzed through the Seurat Package in R (Stuart et al., 2019) keeping only cells having between 1,500 and 30,000 counts and less than 30% of genes coming from the mitochondrial genome. SCTransform was used as the scaling method (Hafemeister & Satija, 2019). The UMAPs were made using 20 dimensions and the clustering resolution was 0.3 for epithelial cells and 1 for enteroendocrine cells.

### Microbiota analysis

Mice were randomly allocated to cages at weaning. Stools were collected either at different time points during mouse lifetime or at sacrifice and stored at −80°C. Metagenome analysis was performed by Eurofins Genomics and SCFA concentration quantification was performed by Creative Proteomics using gas chromatography. To analyze the relative prevalence of *Turicibacter sanguinis* by qPCR in the fecal microbiota, DNA was extracted from the stool after digestion with proteinase K (Sigma Aldrich), followed by isopropanol and ethanol precipitation. 25 ng of DNA were used to perform qPCR to quantify the relative amount of *Turicibacter sanguinis* compared to universal 16S rDNA levels.

### Statistical analysis

Statistical analyses were performed with Graph Pad Prism version 9. All experimental data are expressed as mean ± SEM. The significance of differences between groups was determined by appropriate parametric or non-parametric tests as described in Figure legends.

### Data availability

The datasets produced in this study are available at GSE229812, GSE229813 and GSE229814.

## Acknowledgments

We are grateful to Sumeet Singh Pal and Elif Sema Eski for helpful discussions on single cell transcriptome analyses and Christine Dubois for FACS experiments. The project was funded by the Belgian “Région de Bruxelles-Capitale – Innoviris” (PHD 118 OLFRINGUT grant), the Funds “Fonds David et Alice Van Buuren”, “Fondation Jaumotte-Demoulin” and “Fondation Héger-Masson” and the non-for-profit organisation “Association Recherche Biomédicale et Diagnostic”.

## Author contributions

GD: study concept and design, acquisition of data, analysis and interpretation of data, statistical analysis, drafting of the ms.

ML, AL, FL: acquisition and analysis of data.

YQ, AV, GV, SH, MP: study concept and design, critical revision of the ms, obtained funding, study supervision.

MIG: study concept and design, acquisition of data, analysis and interpretation of data, drafting of the ms, critical revision of the ms, obtained funding, study supervision.

All authors read and approved the final paper.

---

**Figure EV1.**
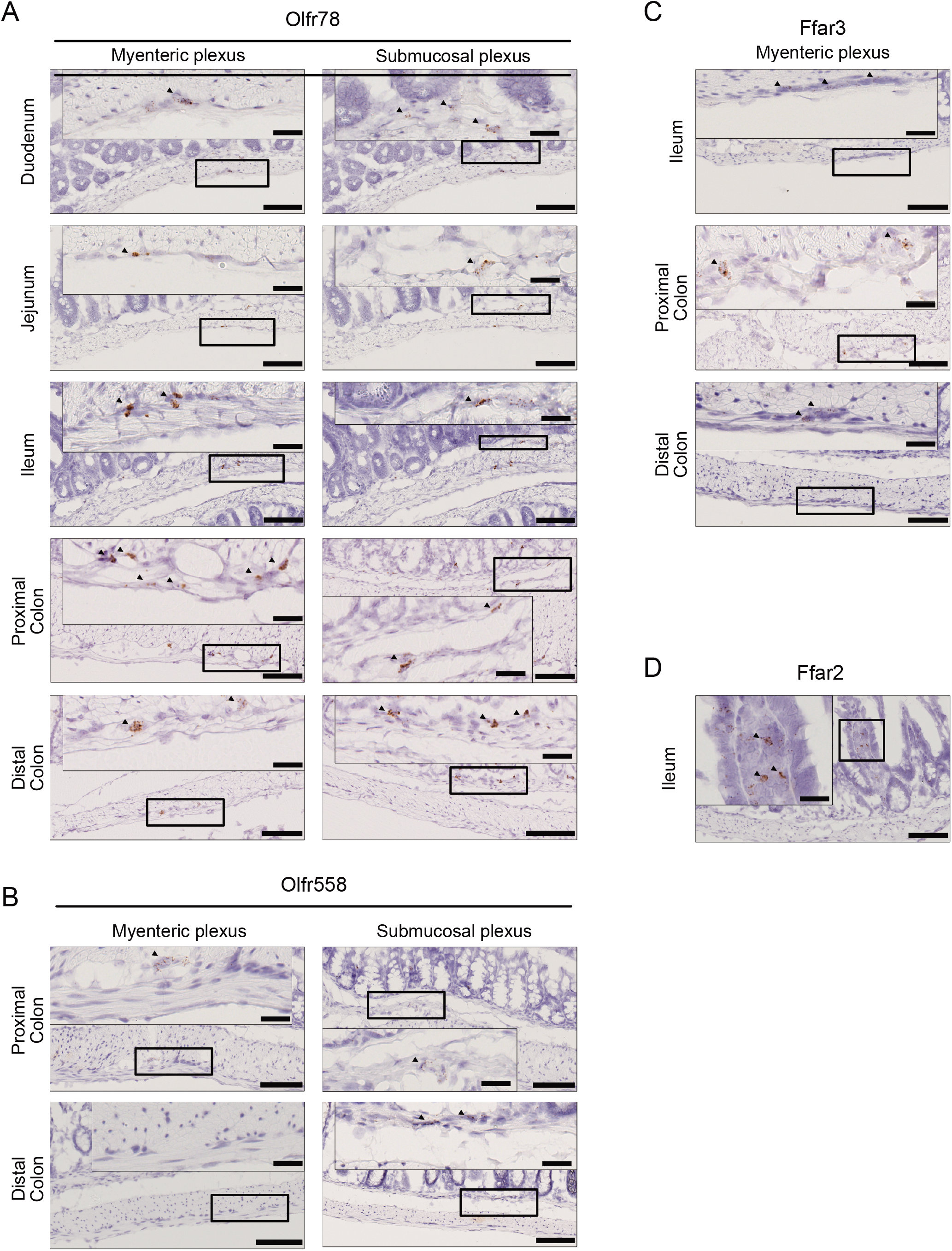
SCFA receptors exhibit unique expression profiles along the small intestine and colon. A. Representative RNAscope pictures of Olfr78 expression in myenteric and submucosal plexuses in the gut. B. Representative RNAscope pictures of Olfr558 expression myenteric and submucosal plexuses in the colon. C. Representative RNAscope pictures of Ffar3 expression in myenteric plexuses in the gut. D. Representative RNAscope pictures of mesenchymal expression of Ffar2 in the Ileum. Data information: scale bars = 25 µm (inset) or 100 µm (low view).

**Figure EV2.**
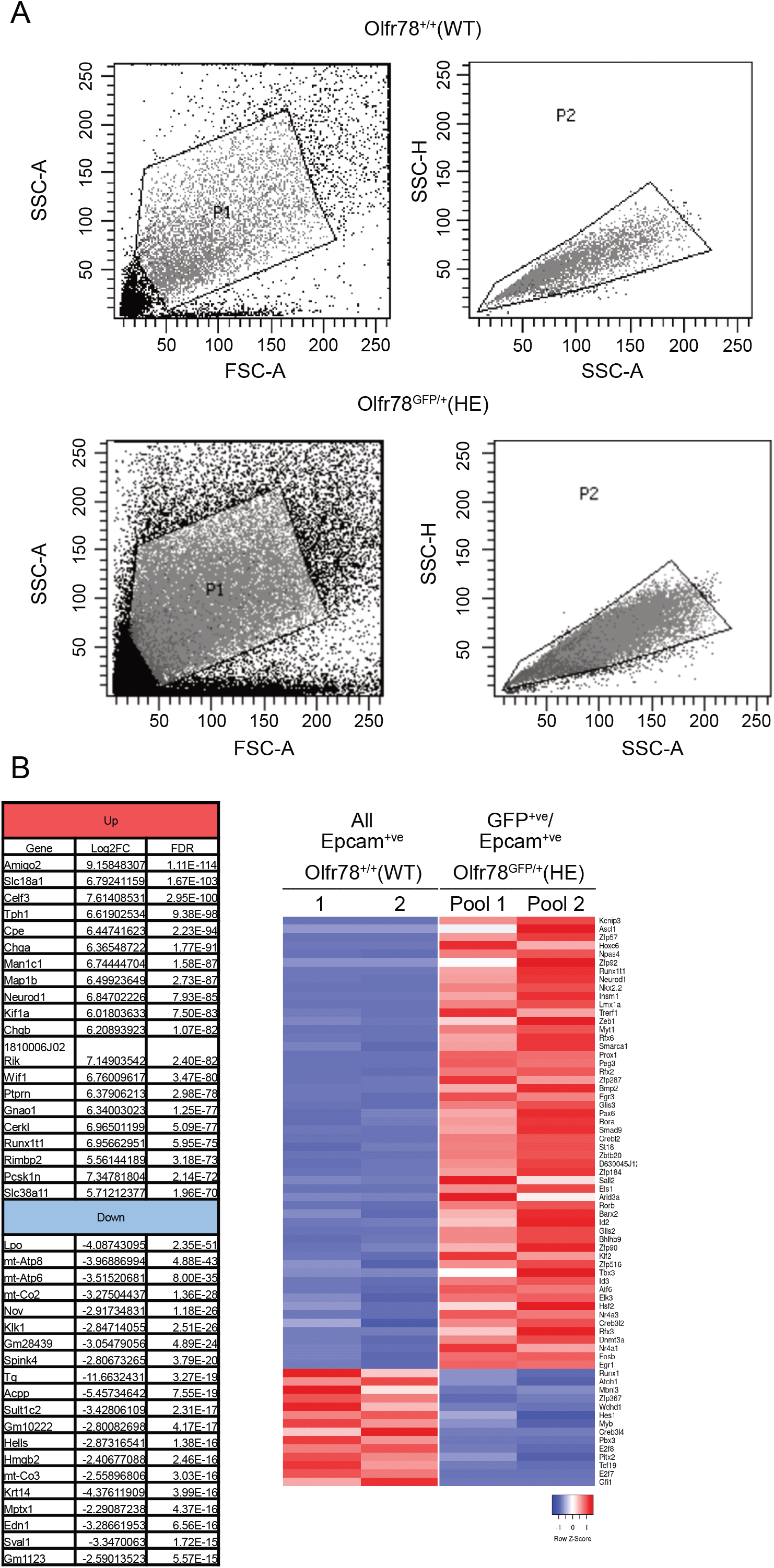
Olfr78 is expressed in different subtypes of enteroendocrine cells in the colon. A. FACS strategy for initial population selection and doublets exclusion. B. Left panel: list of the 20 most up- or downregulated genes in Epcam^+ve^/GFP^+ve^ cells compared to Epcam^+ve^ cells, ranked by FDR. Right panel: list of significantly up and downregulated transcription factors in Epcam^+ve^/GFP^+ve^ cells compared to Epcam^+ve^ cells.

**Figure EV3.**
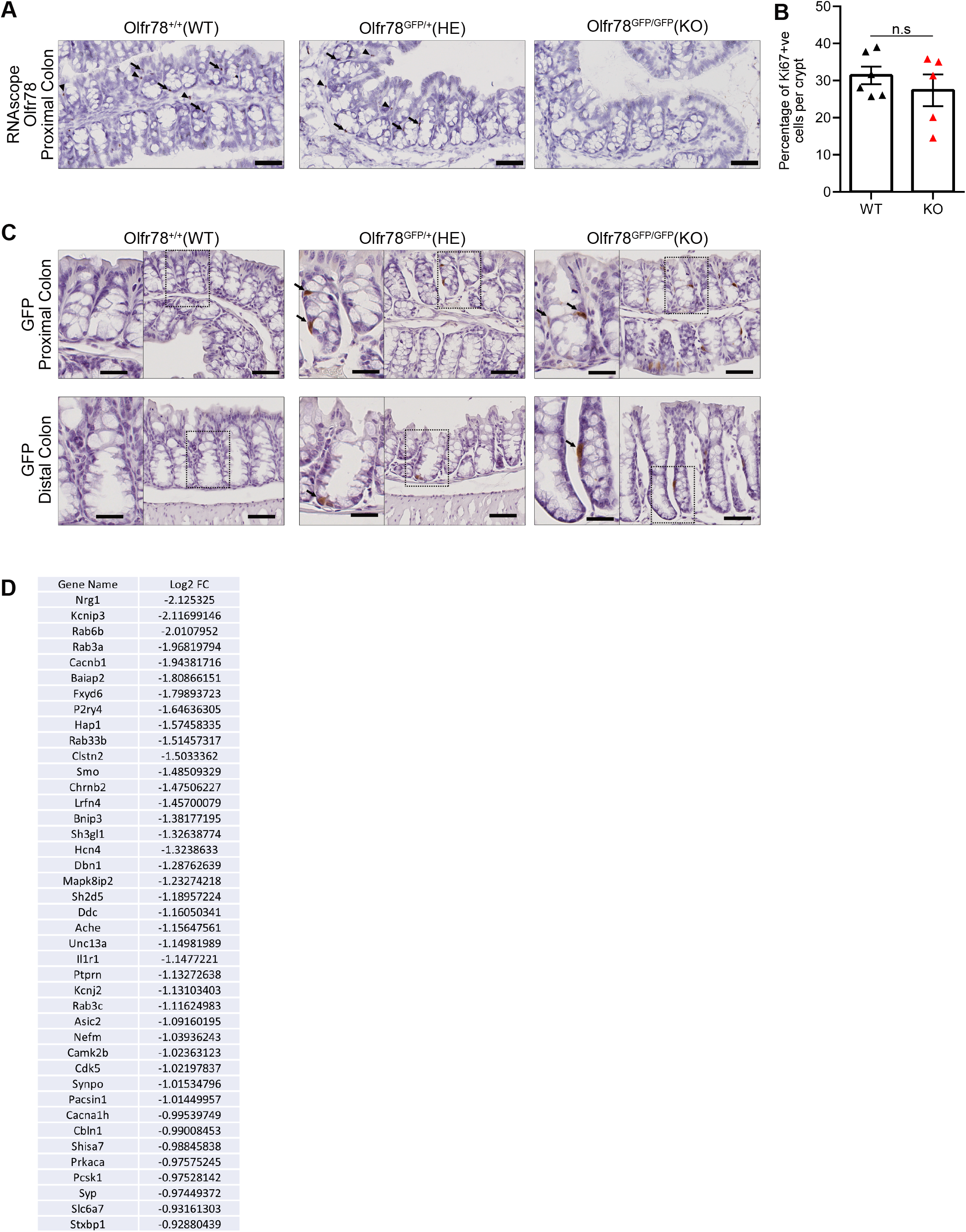
Loss of Olfr78 impairs terminal differentiation into enterochromaffin cells. A. Olfr78 expression in proximal colon of WT, HE and Olfr78-GFP KO mice analyzed by RNAscope. Arrows indicate epithelial cells; arrowheads indicate mesenchymal cells. B. Quantification of Ki67+ve cells in proximal colon of WT and KO mice. Each symbol indicates the value for a given mouse. C. GFP expression in proximal colon of WT, HE and Olfr78-GFP KO mice analyzed by Immunohistochemistry. Arrows indicate epithelial cells. D. List of downregulated genes in KO Epcam^+ve^/GFP^+ve^ cells related to GSEA pre- or post-synapse gene lists, ranked by Log_2_(Fold Change). Data information: scale bars = 25 µm (inset) or 50 µm (low view). Data are represented as mean ± SEM.; n.s = not significant; Mann Whitney (B).

**Figure EV4.**
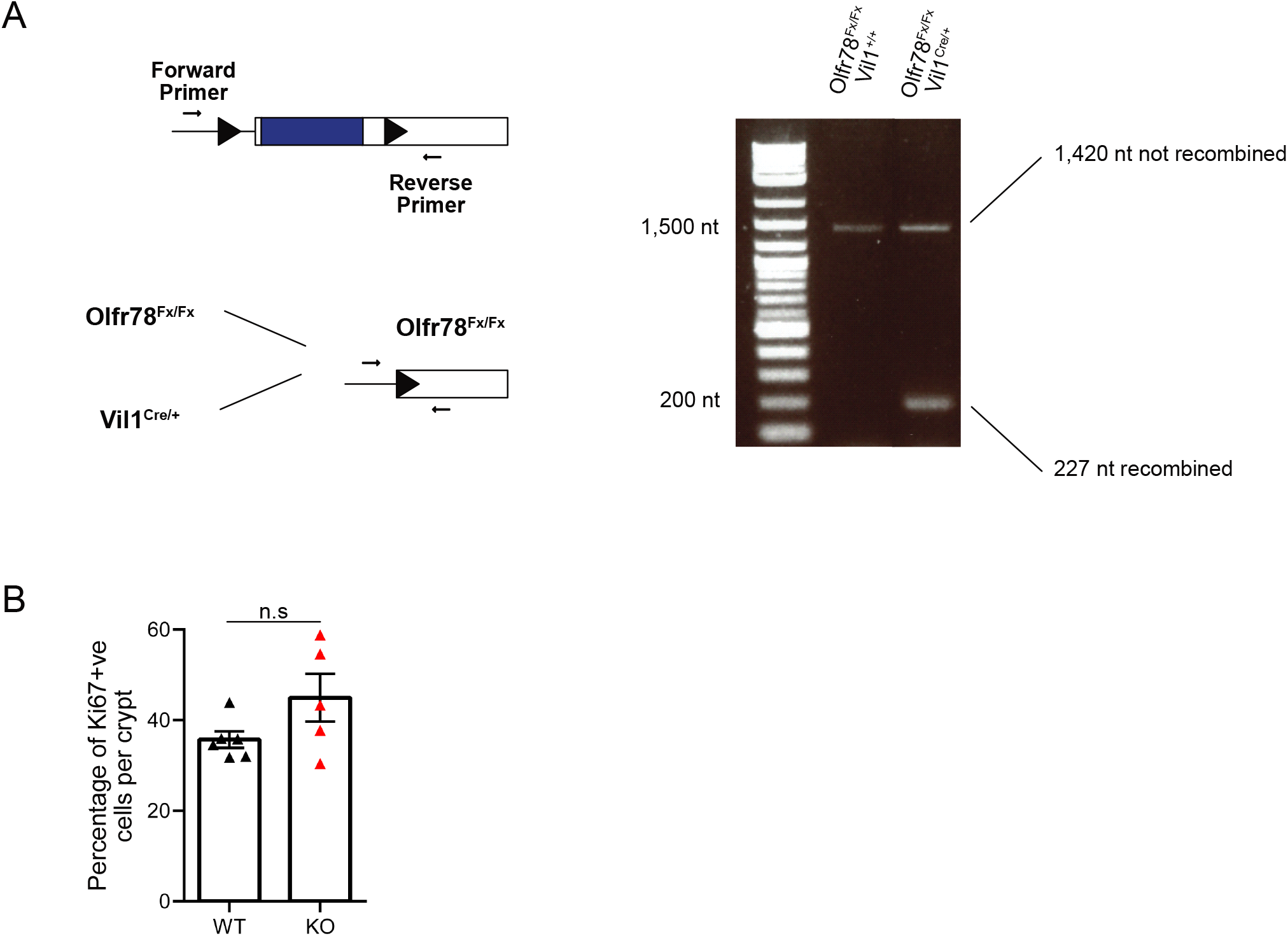
Terminal differentiation into serotonin-producing cells is regulated by epithelial Olfr78 expression. A. Left: PCR strategy for loxP sites recombination verification in Olfr78^Fx/Fx^-Vil1^Cre/WT^. Right: Gel electrophoresis showing WT and recombinant bands in Olfr78^Fx/Fx^-Vil1^+/+^. nt = nucleotide. B. Quantification of Ki67+ve cells in proximal colon of WT and KO mice. Each symbol indicates the value for a given mouse. Data information: Data are represented as mean ± SEM.; n.s = not significant; Mann Whitney (B).

**Figure EV5.**
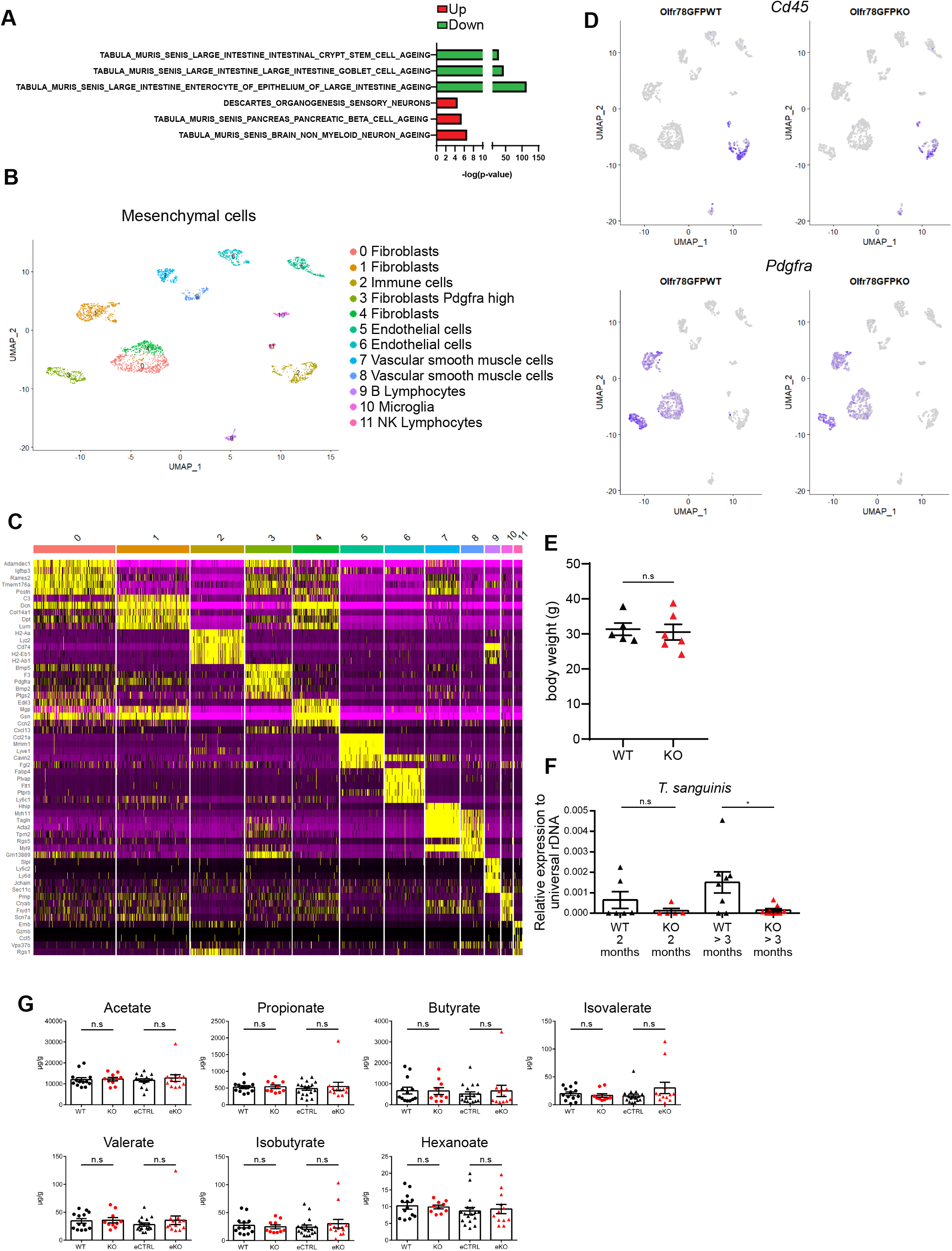
Loss of Olfr78 expression alters colon homeostasis. A. Modulated Mol-Sig GSEA C8 cell signature in the transcriptome of Olfr78-GFP KO vs Olfr78- GFP WT crypts. B. ScRNAsequencing-based UMAP of mesenchymal cells from one WT and one Olfr78-GFP KO mice after merging (n= 1,505 for the WT, n= 1,514 for the KO). C. Top 5 markers of each cluster of the UMAP in B. D. Expression of *Cd45* and *Pdgfra* in WT and KO cells split from the UMAP in B. E. Weight of adult Olfr78-GFP mice. Each symbol indicates the value for a given mouse. F. Analysis of *Turicibacter sanguinis* prevalence by qPCR in the fecal microbiota of Olfr78-GFP mice at different ages. Each symbol indicates the value for a given mouse. G. Quantification of fecal SCFA concentrations. Each symbol indicates the value for a given mouse. Data information: Data are represented as mean ± SEM.; n.s = not significant; * P <0.05, Mann whitney (E, F, G).

## REFERENCES

Abaffy, T., Bain, J. R., Muehlbauer, M. J., Spasojevic, I., Lodha, S., Bruguera, E., O’Neal, S. K., Kim, S. Y., & Matsunami, H. (2018). A testosterone metabolite 19-hydroxyandrostenedione induces neuroendocrine trans-differentiation of prostate cancer cells via an ectopic olfactory receptor. Frontiers in Oncology, 8(MAY). https://doi.org/10.3389/fonc.2018.00162

Asle, B. W., Turvey, M., Larina, O., Thorn, P., Skepper, J., Morton, A. J., & Edwardson, J. M. (2005). Syncollin is required for efficient zymogen granule exocytosis. In Biochem. J (Vol. 385).

Audouze, K., Tromelin, A., Le Bon, A. M., Belloir, C., Petersen, R. K., Kristiansen, K., Brunak, S., & Taboureau, O. (2014). Identification of odorant-receptor interactions by global mapping of the human odorome. PLoS ONE, 9(4). https://doi.org/10.1371/journal.pone.0093037

Basak, O., Beumer, J., Wiebrands, K., Seno, H., van Oudenaarden, A., & Clevers, H. (2017). Induced Quiescence of Lgr5+ Stem Cells in Intestinal Organoids Enables Differentiation of Hormone-Producing Enteroendocrine Cells. Cell Stem Cell, 20(2), 177–190.e4. https://doi.org/10.1016/j.stem.2016.11.001

Bellono, N. W., Bayrer, J. R., Leitch, D. B., Castro, J., Zhang, C., O’Donnell, T. A., Brierley, S. M., Ingraham, H. A., & Julius, D. (2017). Enterochromaffin Cells Are Gut Chemosensors that Couple to Sensory Neural Pathways. Cell, 170(1), 185–198.e16. https://doi.org/10.1016/j.cell.2017.05.034

Beumer, J., Artegiani, B., Post, Y., Reimann, F., Gribble, F., Nguyen, T. N., Zeng, H., Van den Born, M., Van Es, J. H., & Clevers, H. (2018). Enteroendocrine cells switch hormone expression along the crypt-to-villus BMP signalling gradient. In Nature Cell Biology (Vol. 20, Issue 8, pp. 909–916). Nature Publishing Group. https://doi.org/10.1038/s41556-018-0143-y

Billing, L. J., Larraufie, P., Lewis, J., Leiter, A., Li, J., Lam, B., Yeo, G. S., Goldspink, D. A., Kay, R. G., Gribble, F. M., & Reimann, F. (2019). Single cell transcriptomic profiling of large intestinal enteroendocrine cells in mice – Identification of selective stimuli for insulin-like peptide-5 and glucagon-like peptide-1 co-expressing cells. Molecular Metabolism, 29, 158–169. https://doi.org/10.1016/j.molmet.2019.09.001

Bohórquez, D. V., Samsa, L. A., Roholt, A., Medicetty, S., Chandra, R., & Liddle, R. A. (2014). An enteroendocrine cell - Enteric glia connection revealed by 3D electron microscopy. PLoS ONE, 9(2). https://doi.org/10.1371/journal.pone.0089881

Bohórquez, D. V., Shahid, R. A., Erdmann, A., Kreger, A. M., Wang, Y., Calakos, N., Wang, F., & Liddle, R. A. (2015). Neuroepithelial circuit formed by innervation of sensory enteroendocrine cells. Journal of Clinical Investigation, 125(2), 782–786. https://doi.org/10.1172/JCI78361

Bozza, T., Vassalli, A., Fuss, S., Zhang, J. J., Weiland, B., Pacifico, R., Feinstein, P., & Mombaerts, P. (2009). Mapping of Class I and Class II Odorant Receptors to Glomerular Domains by Two Distinct Types of Olfactory Sensory Neurons in the Mouse. Neuron, 61(2), 220–233. https://doi.org/10.1016/j.neuron.2008.11.010

Brown, D., Sorscher, E. J., Ausiello, D. A., & Benos, D. J. (1989). Immunocytochemical localization of Na+ channels in rat kidney medulla. www.physiology.org/journal/ajprenal

Buck, L., & Axel, R. (1991). A Novel Multigene Family May Encode Odorant Receptors: A Molecular Basis for Odor Recognition. In Cell (Vol. 65).

Chen, C., & Lodish, H. F. (2014). Global analysis of induced transcription factors and cofactors identifies Tfdp2 as an essential coregulator during terminal erythropoiesis. Experimental Hematology, 42(6), 464–476.e5. https://doi.org/10.1016/j.exphem.2014.03.001

Christiansen, C. B., Buur, M., Gabe, N., Svendsen, B., Dragsted, L. O., Rosenkilde, M. M., & Holst, J. J. (2018). The impact of short-chain fatty acids on GLP-1 and PYY secretion from the isolated perfused rat colon. Am J Physiol Gastrointest Liver Physiol, 315, 53–65. https://doi.org/10.1152/ajpgi.00346.2017.-The

Cong, J., Zhou, P., & Zhang, R. (2022). Intestinal Microbiota-Derived Short Chain Fatty Acids in Host Health and Disease. In Nutrients (Vol. 14, Issue 9). MDPI. https://doi.org/10.3390/nu14091977

Egerod, K. L., Engelstoft, M. S., Grunddal, K. V., Nøhr, M. K., Secher, A., Sakata, I., Pedersen, J., Windeløv, J. A., Füchtbauer, E. M., Olsen, J., Sundler, F., Christensen, J. P., Wierup, N., Olsen, J. V., Holst, J. J., Zigman, J. M., Poulsen, S. S., & Schwartz, T. W. (2012). A major lineage of enteroendocrine cells coexpress CCK, secretin, GIP, GLP-1, PYY, and neurotensin but not somatostatin. Endocrinology, 153(12), 5782–5795. https://doi.org/10.1210/en.2012-1595

Fleischer, J., Bumbalo, R., Bautze, V., Strotmann, J., & Breer, H. (2015). Expression of odorant receptor Olfr78 in enteroendocrine cells of the colon. Cell and Tissue Research, 361(3), 697– 710. https://doi.org/10.1007/s00441-015-2165-0

Fung, T. C., Vuong, H. E., Luna, C. D. G., Pronovost, G. N., Aleksandrova, A. A., Riley, N. G., Vavilina, A., McGinn, J., Rendon, T., Forrest, L. R., & Hsiao, E. Y. (2019). Intestinal serotonin and fluoxetine exposure modulate bacterial colonization in the gut. In Nature Microbiology (Vol. 4, Issue 12, pp. 2064–2073). Nature Research. https://doi.org/10.1038/s41564-019-0540-4

Gehart, H., van Es, J. H., Hamer, K., Beumer, J., Kretzschmar, K., Dekkers, J. F., Rios, A., & Clevers, H. (2019). Identification of Enteroendocrine Regulators by Real-Time Single-Cell Differentiation Mapping. Cell, 176(5), 1158–1173.e16. https://doi.org/10.1016/j.cell.2018.12.029

Gribble, F. M., & Reimann, F. (2019). Function and mechanisms of enteroendocrine cells and gut hormones in metabolism. In Nature Reviews Endocrinology (Vol. 15, Issue 4, pp. 226–237). Nature Publishing Group. https://doi.org/10.1038/s41574-019-0168-8

Haber, A. L., Biton, M., Rogel, N., Herbst, R. H., Shekhar, K., Smillie, C., Burgin, G., Delorey, T. M., Howitt, M. R., Katz, Y., Tirosh, I., Beyaz, S., Dionne, D., Zhang, M., Raychowdhury, R., Garrett, W. S., Rozenblatt-Rosen, O., Shi, H. N., Yilmaz, O., … Regev, A. (2017). A single-cell survey of the small intestinal epithelium. Nature, 551(7680), 333–339. https://doi.org/10.1038/nature24489

Habib, A. M., Richards, P., Cairns, L. S., Rogers, G. J., Bannon, C. A. M., Parker, H. E., Morley, T. C. E., Yeo, G. S. H., Reimann, F., & Gribble, F. M. (2012). Overlap of endocrine hormone expression in the mouse intestine revealed by transcriptional profiling and flow cytometry. Endocrinology, 153(7), 3054–3065. https://doi.org/10.1210/en.2011-2170

Hafemeister, C., & Satija, R. (2019). Normalization and variance stabilization of single-cell RNA-seq data using regularized negative binomial regression. Genome Biology, 20(1). https://doi.org/10.1186/s13059-019-1874-1

Kim, M. H., Kang, S. G., Park, J. H., Yanagisawa, M., & Kim, C. H. (2013). Short-chain fatty acids activate GPR41 and GPR43 on intestinal epithelial cells to promote inflammatory responses in mice. Gastroenterology, 145(2). https://doi.org/10.1053/j.gastro.2013.04.056

Kotlo, K., Anbazhagan, A. N., Priyamvada, S., Jayawardena, D., Kumar, A., Chen, Y., Xia, Y., Finn, P. W., Perkins, D. L., Dudeja, P. K., & Layden, B. T. (2020). The olfactory G protein-coupled receptor (Olfr-78/OR51E2) modulates the intestinal response to colitis. Am J Physiol Cell Physiol, 318, 502–513. https://doi.org/10.1152/ajpcell.00454.2019.-Olfactory

Kuo, C. S., Darmanis, S., de Arce, A. D., Liu, Y., Almanzar, N., Wu, T. T. H., Quake, S. R., & Krasnow, M. A. (2022). Neuroendocrinology of the lung revealed by single-cell RNA sequencing. ELife, 11. https://doi.org/10.7554/ELIFE.78216

Le Poul, E., Loison, C., Struyf, S., Springael, J. Y., Lannoy, V., Decobecq, M. E., Brezillon, S., Dupriez, V., Vassart, G., Van Damme, J., Parmentier, M., & Detheux, M. (2003). Functional characterization of human receptors for short chain fatty acids and their role in polymorphonuclear cell activation. Journal of Biological Chemistry, 278(28), 25481–25489. https://doi.org/10.1074/jbc.M301403200

Lee, S.-J., Depoortere, I., & Hatt, H. (2019). Therapeutic potential of ectopic olfactory and taste receptors. Nature Reviews Drug Discovery, 18, 116–138. https://doi.org/10.1038/s41573

Lund, M. L., Egerod, K. L., Engelstoft, M. S., Dmytriyeva, O., Theodorsson, E., Patel, B. A., & Schwartz, T. W. (2018). Enterochromaffin 5-HT cells – A major target for GLP-1 and gut microbial metabolites. Molecular Metabolism, 11, 70–83. https://doi.org/10.1016/j.molmet.2018.03.004

Nishida, A., Miyamoto, J., Shimizu, H., & Kimura, I. (2021). Gut microbial short-chain fatty acids-mediated olfactory receptor 78 stimulation promotes anorexigenic gut hormone peptide YY secretion in mice. Biochemical and Biophysical Research Communications, 557, 48–54. https://doi.org/10.1016/j.bbrc.2021.03.167

Nøhr, M. K., Pedersen, M. H., Gille, A., Egerod, K. L., Engelstoft, M. S., Husted, A. S., Sichlau, R. M., Grunddal, K. V., Poulsen, S. S., Han, S., Jones, R. M., Offermanns, S., & Schwartz, T. W. (2013). GPR41/FFAR3 and GPR43/FFAR2 as cosensors for short-chain fatty acids in enteroendocrine cells vs FFAR3 in enteric neurons and FFAR2 in enteric leukocytes. Endocrinology, 154(10), 3552–3564. https://doi.org/10.1210/en.2013-1142

Perland, E., Hellsten, S. V., Schweizer, N., Arapi, V., Rezayee, F., Bushra, M., & Fredriksson, R. (2017). Structural prediction of two novel human atypical SLC transporters, MFSD4A and MFSD9, and their neuroanatomical distribution in mice. PLoS ONE, 12(10). https://doi.org/10.1371/journal.pone.0186325

Piccand, J., Vagne, C., Blot, F., Meunier, A., Beucher, A., Strasser, P., Lund, M. L., Ghimire, S., Nivlet, L., Lapp, C., Petersen, N., Engelstoft, M. S., Thibault-Carpentier, C., Keime, C., Correa, S. J., Schreiber, V., Molina, N., Schwartz, T. W., De Arcangelis, A., & Gradwohl, G. (2019). Rfx6 promotes the differentiation of peptide-secreting enteroendocrine cells while repressing genetic programs controlling serotonin production. Molecular Metabolism, 29, 24–39. https://doi.org/10.1016/j.molmet.2019.08.007

Pluznick, J. L. (2016). Gut microbiota in renal physiology: focus on short-chain fatty acids and their receptors. In Kidney International (Vol. 90, Issue 6, pp. 1191–1198). Elsevier B.V. https://doi.org/10.1016/j.kint.2016.06.033

Pluznick, J. L., Protzko, R. J., Gevorgyan, H., Peterlin, Z., Sipos, A., Han, J., Brunet, I., Wan, L. X., Rey, F., Wang, T., Firestein, S. J., Yanagisawa, M., Gordon, J. I., Eichmann, A., Peti-Peterdi, J., & Caplan, M. J. (2013). Olfactory receptor responding to gut microbiotaderived signals plays a role in renin secretion and blood pressure regulation. Proceedings of the National Academy of Sciences of the United States of America, 110(11), 4410–4415. https://doi.org/10.1073/pnas.1215927110

Psichas, A., Sleeth, M. L., Murphy, K. G., Brooks, L., Bewick, G. A., Hanyaloglu, A. C., Ghatei, M. A., Bloom, S. R., & Frost, G. (2015). The short chain fatty acid propionate stimulates GLP-1 and PYY secretion via free fatty acid receptor 2 in rodents. International Journal of Obesity, 39(3), 424–429. https://doi.org/10.1038/ijo.2014.153

Roth, K. A., Kim, S., & Gordon, J. I. (1992). Immunocytochemical studies suggest two pathways for enteroendocrine cell differentiation in the colon. www.physiology.org/journal/ajpgi

Saito, H., Chi, Q., Zhuang, H., Matsunami, H., & Mainland, J. D. (2009). Odor Coding by a Mammalian Receptor Repertoire. Science Signaling. https://doi.org/10.1126/scisignal

Sato, T., Stange, D. E., Ferrante, M., Vries, R. G. J., Van Es, J. H., Van Den Brink, S., Van Houdt, W. J., Pronk, A., Van Gorp, J., Siersema, P. D., & Clevers, H. (2011). Long-term expansion of epithelial organoids from human colon, adenoma, adenocarcinoma, and Barrett’s epithelium. Gastroenterology, 141(5), 1762–1772. https://doi.org/10.1053/j.gastro.2011.07.050

Serizawa, S., Miyamichi, K., & Sakano, H. (2004). One neuron-one receptor rule in the mouse olfactory system. In Trends in Genetics (Vol. 20, Issue 12, pp. 648–653). https://doi.org/10.1016/j.tig.2004.09.006

Stuart, T., Butler, A., Hoffman, P., Hafemeister, C., Papalexi, E., Mauck, W. M., Hao, Y., Stoeckius, M., Smibert, P., & Satija, R. (2019). Comprehensive Integration of Single-Cell Data. Cell, 177(7), 1888–1902.e21. https://doi.org/10.1016/j.cell.2019.05.031

Subramanian, A., Tamayo, P., Mootha, V. K., Mukherjee, S., Ebert, B. L., Gillette, M. A., Paulovich, A., Pomeroy, S. L., Golub, T. R., Lander, E. S., & Mesirov, J. P. (2005). Gene set enrichment analysis: A knowledge-based approach for interpreting genome-wide expression profiles. www.pnas.orgcgidoi10.1073pnas.0506580102

Tolhurst, G., Heffron, H., Lam, Y. S., Parker, H. E., Habib, A. M., Diakogiannaki, E., Cameron, J., Grosse, J., Reimann, F., & Gribble, F. M. (2012). Short-chain fatty acids stimulate glucagon-like peptide-1 secretion via the G-protein-coupled receptor FFAR2. Diabetes, 61(2), 364–371. https://doi.org/10.2337/db11-1019

Tsuruta, T., Saito, S., Osaki, Y., Hamada, A., Aoki-Yoshida, A., & Sonoyama, K. (2016). Organoids as an ex vivo model for studying the serotonin system in the murine small intestine and colon epithelium. Biochemical and Biophysical Research Communications, 474(1), 161–167. https://doi.org/10.1016/j.bbrc.2016.03.165

Wang, Y. C., Zuraek, M. B., Kosaka, Y., Ota, Y., German, M. S., Deneris, E. S., Bergsland, E. K., Donner, D. B., Warren, R. S., & Nakakura, E. K. (2010). The ETS oncogene family transcription factor FEV identifies serotonin-producing cells in normal and neoplastic small intestine. Endocrine-Related Cancer, 17(1), 283–291. https://doi.org/10.1677/ERC-09-0243

Yadav, G. P., Zheng, H., Yang, Q., Douma, L. G., Bloom, L. B., & Jiang, Q. X. (2018). Secretory granule protein chromogranin B (CHGB) forms an anion channel in membranes. Life Science Alliance, 1(5). https://doi.org/10.26508/lsa.201800139

